# De Novo Protocell Membrane Formation Fueled by Primitive Metabolites

**DOI:** 10.1101/2025.05.10.653202

**Authors:** Satyam Khanal, Alessandro Fracassi, Alexander Harjung, Michael D. Burkart, Neal K. Devaraj

## Abstract

Lipid membranes define cell boundaries, acting as gatekeepers for transport and signaling. A central principle in biology is that all cellular membranes descend from a common ancestral membrane, as they cannot be generated in the absence of preexisting lipid structures. It is thus unclear how the first protocell membranes originated from membrane-less precursors. Here we demonstrate the de novo generation of lipid bilayers in the absence of any preexisting membranes, membrane-bound proteins, or lipid nanostructure templates. Using acetate as a two-carbon precursor, lipid tails are constructed by soluble enzymes and spontaneously conjugate to cysteine backbones, forming diacyl lipids that assemble into vesicles. Pore-forming peptides facilitate precursor transport into vesicles, allowing the continuous generation of new lipids. Formation of glycolipid membranes creates compartments that can maintain proton gradients. The de novo formation of membrane bilayers using primitive chemical building blocks may have been an intermediary step between primitive prebiotic biochemistry and the emergence of cellular life.

## Introduction

Cells compartmentalize themselves from their environment through bilayer membranes composed primarily of polar lipids such as phospholipids and glycolipids. Membrane lipid biosynthesis requires the action of several enzymes, including many that are membrane bound (Figure 1a) (*1*). The requirement of membrane bound proteins for generating the lipids that constitute the membranes themselves has led to speculation that membranes, like genetic material, are inherited, and that all membranes can be traced back to a common ancestral membrane (*2, 3*). While this might be satisfactory for understanding how cellular membranes propagate in extant organisms, membrane heredity presents a paradox regarding how the first membranes arose de novo, i.e., in the absence of preexisting membranes. How could lipid membranes have been present without membrane proteins, such as acyltransferases for phospholipid synthesis or translocons for membrane protein synthesis? The spontaneous formation of membranes from primordial broths that supported primitive biochemistry has long been proposed (*3*), but no experiments have yet led to the generation of lipid membranes purely from structurally simple metabolic precursors. Several reports have demonstrated activity from reconstituting lipid membrane biogenesis pathways in artificial cells, but the lipid yields are typically low; and de novo membrane formation is not feasible, as the key enzymes in lipid biosynthesis are themselves membrane bound (*4–6*). Previous investigations have also demonstrated chemoenzymatic membrane formation from reactive surfactants, but these studies require the use of complex, long-chain hydrocarbon lipid precursors that form lipid nanoassemblies to template membrane formation (*7, 8*).

**Figure 1.**
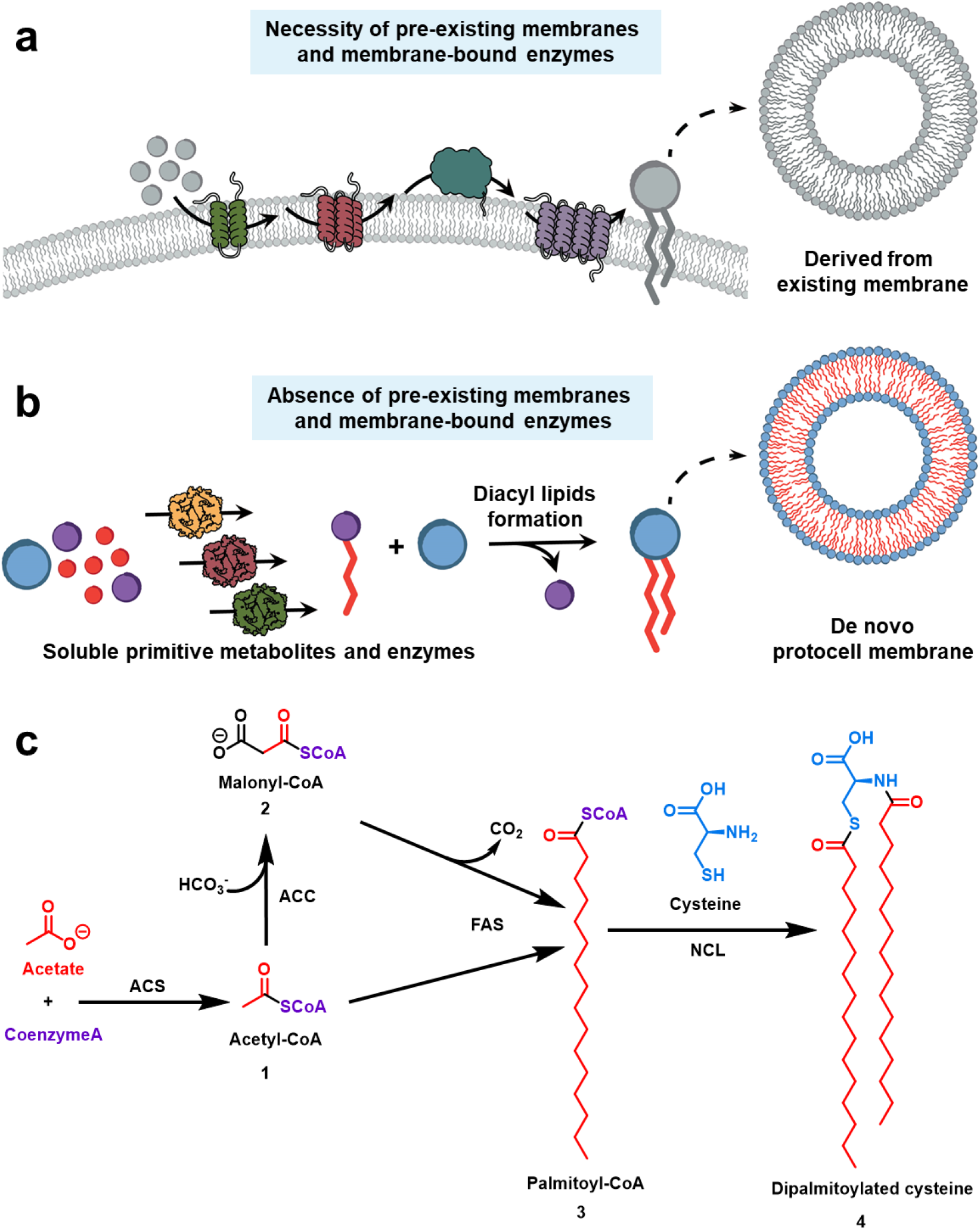
Schematic representation of de novo membrane formation driven by lipid synthesis from primitive metabolites. **(a)** Representative illustration of a generalized lipid biosynthesis process leading to the formation of new lipid membranes. The process depends on the presence of pre-existing membranes and membrane-bound proteins necessary for lipid synthesis. **(b)** Proposed approach for the formation of de novo generated membranes. Soluble primitive metabolites are converted into reactive single chain amphiphiles by soluble enzymes, which react with primitive headgroup precursors to produce diacylated lipids that self-assemble into membranes. Membranes are formed in the absence of any pre-existing membranes, long-chain hydrocarbon lipids, or transmembrane proteins. **(c)** Reaction scheme for the synthesis of lipid **4**. Three enzymes, ACS, ACC, and FAS mediate the formation of palmitoyl-CoA **3** using acetate as a carbon source. Palmitoyl-CoA chemoselectively diacylates cysteine to generate dipalmitoylated cysteine lipid **4**, which is capable of spontaneous self-assembly, forming bilayer vesicles.

Here we demonstrate de novo formation of protocell membranes from primitive metabolites using a minimal set of soluble proteins. By generating acyl precursors using soluble enzymes and replacing the steps normally catalyzed by membrane-bound acyltransferases with chemical steps, simplified analogs of biological membrane lipids can be fully synthesized in the absence of preexisting membranes (Figure 1b). Acyl chains are enzymatically generated starting from acetate, a structurally simple two-carbon precursor and universal metabolite that can be formed abiotically through geochemical processes (*9–11*). In situ generated acyl chains spontaneously diacylate cysteine, an essential amino-acid that has recently been shown to be important in prebiotic metabolism as a catalyst for peptide bond formation (*12*). The diacylation products are polar lipids that resemble natural diacyl membrane lipids and spontaneously assemble to form protocell vesicles (Figure 1c). Short pore-forming peptides can spontaneously insert into the de novo formed vesicles, forming channels that permit the transport of new precursors for continual membrane synthesis using entrapped proteins. Adapting the approach with more complex polar headgroups enables the formation of diacyl glycolipids, which are non-ionizable and self-assemble to form membranes that can maintain proton gradients, a conserved feature found in all life on Earth. Our work demonstrates for the first time that de novo membrane generation is feasible, establishing that cell-like membranes using simple carbon feedstocks to fuel membrane synthesis could have been a viable strategy for the formation and propagation of early cells. Such simplified membrane formation may have served as an intermediate evolutionary step, bridging simple prebiotic biochemistry and the complex lipid membranes that are an essential feature of all extant organisms (*13*).

## Results and Discussion

To form lipid membranes de novo from non-lipidic metabolites, the long-chain hydrocarbon tails that are essential for membrane assembly must be synthesized from structurally simple, soluble molecules. Starting from acetate, we combined three enzymatic reactions to build the long-chain acyl-CoA thioester palmitoyl-CoA. We used *E. coli* acetyl-CoA synthetase (ACS) to convert acetate to acetyl CoA (*14*), human acetyl-CoA carboxylase (ACC) for producing malonyl-CoA from the generated acetyl-CoA (*15, 16*), and a type I fatty acid synthase from *Corynebacterium glutamicum* (FAS) to synthesize palmitoyl-CoA from acetyl-CoA and malonyl-CoA (Figure 1c) (*17*). Initially, we individually assessed the activity of ACS, ACC, and FAS, which provided yields of 66% for acetyl-CoA, 71% for malonyl-CoA, and 63% for palmitoyl-CoA, over 4 hours (Figure S1-S3). Combining the three enzymes provided palmitoyl-CoA with a 42% yield within 4 hours (Figure 2a, and S4). No palmitoyl-CoA was observed in control experiments lacking any of the three enzymes or in the absence of acetate (Figure 2a).

**Figure 2.**
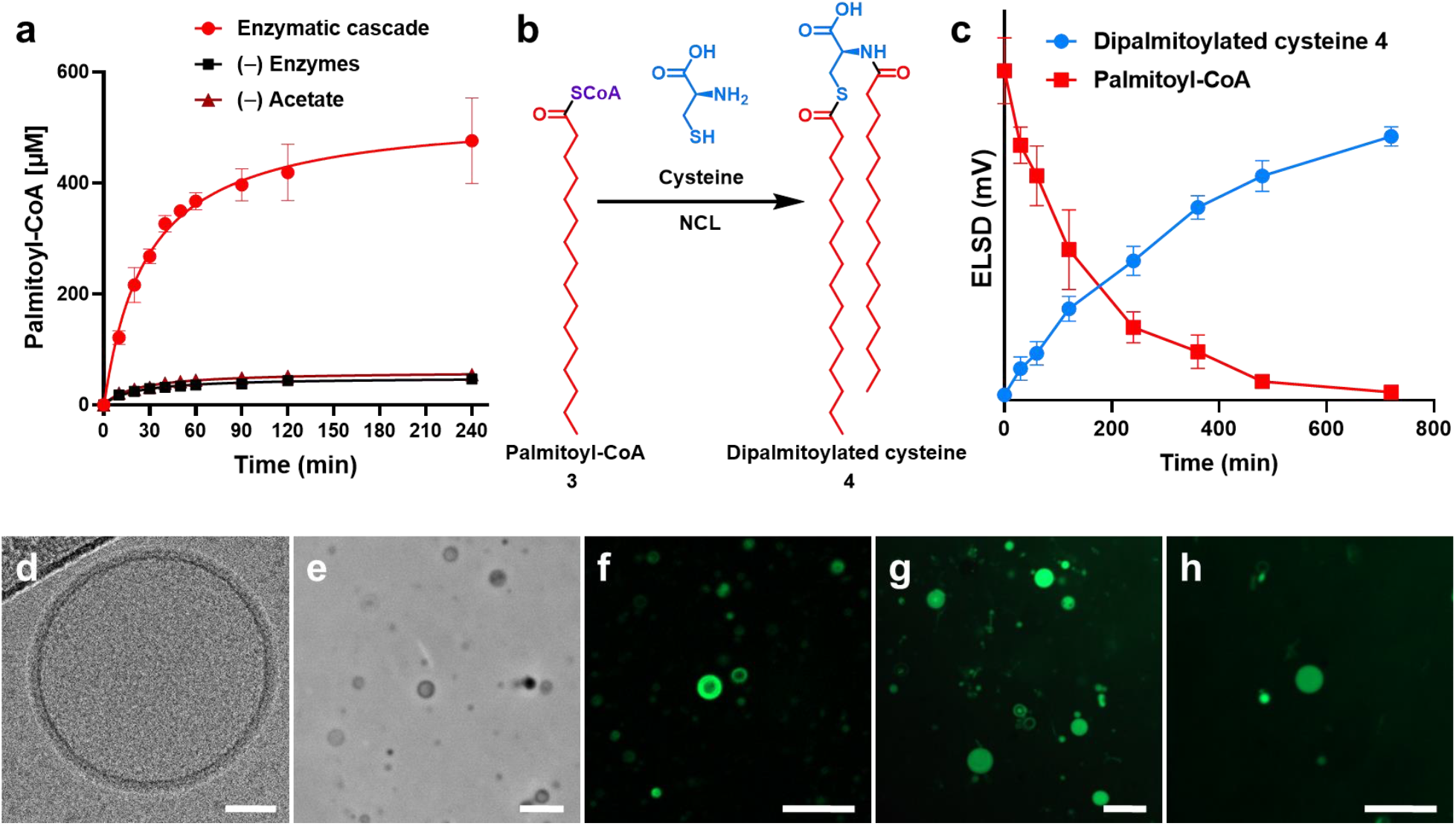
Enzymatic generation of palmitoyl-CoA from acetate, chemical synthesis of dipalmitoylated cysteine 4, and its spontaneous self-assembly into vesicles. **(a)** Palmitoyl-CoA synthesis from acetate mediated by the enzymatic cascade of ACS, ACC, and FAS. Data points presented as means ± s.d. (*n* = 3 technically independent samples). **(b)** Reaction scheme of dipalmitoylated cysteine **4** synthesis from palmitoyl-CoA and cysteine in bicine buffer pH 8.5. **(c)** Formation of lipid **4** over 12 h (blue) and concomitant consumption of palmitoyl-CoA (red) in bicine buffer pH 8.5. Data points presented as means ± s.d. (*n* = 3 technically independent samples). **(d)** Cryo-EM image of a vesicle of lipid **4** generated in situ from the reaction between palmitoyl-CoA and cysteine. Scale bar = 20 nm. **(e)** Phase-contrast microscopy image of membrane-bound vesicles resulting from the synthesis of lipid **4** and its spontaneous self-assembly in the presence of 10 mol% cholesterol. Scale bar = 5 µm. **(f)** Fluorescence microscopy image of vesicles of **4** stained with 0.1 mol% BODIPY-FL. Scale bar = 5 µm. **(g)** Fluorescence microscopy image of vesicles of **4** showing encapsulation of HPTS in the aqueous core. Scale bar = 5 µm. **(h)** Fluorescence microscopy image of vesicles of **4** showing encapsulation of GFP in the aqueous core. Scale bar = 5 µm.

In living cells, long-chain acyl-CoA thioesters like palmitoyl-CoA are used to acylate hydrophilic head groups and generate polar lipids like phospholipids, sphingolipids, and glycolipids. The acylation reactions are catalyzed by proteins which typically require integration into lipid membranes to be fully functional (*18–20*). We previously demonstrated that thioesters can spontaneously acylate aminothiol functional groups by native chemical ligation (NCL) (*21–23*). Recent work has observed that protocells can be generated by the spontaneous dual *N*- and *S*-acylation of cysteine (*24*). However, pre-synthetized unnatural long-chain thioesters were required to template membrane formation, and true de novo synthesis was not demonstrated. We hypothesized that cysteine might be similarly diacylated in the presence of in situ generated palmitoyl-CoA thioesters forming a diacyl lipid capable of assembling into membranes. While cells membranes are often formed by diacylation of a glycerol backbone, it has recently been discovered that certain thermophilic bacteria form a large percentage of their membrane lipids by diacylation of a serine amino acid derived backbone (*25, 26*). Amino acid diacylation may be a biochemically relevant mechanism for forming membrane lipids and such lipids could have served as primordial precursors to cellular lipids.

We initially tested whether palmitoyl-CoA was capable of diacylating cysteine. We mixed 2 mM of cysteine with 4 mM palmitoyl-CoA at 37 °C in bicine buffer at pH 8.5 (Figure 2b) and observed significant dipalmitoylated cysteine **4** formation by HPLC-ELSD-MS, along with the consumption of palmitoyl-CoA (Figure 2c). We confirmed synthesis of **4** by comparison with a chemically synthesized standard (Figures S5). After 8 hours, we observed 1.6±0.3 mM of lipid **4**, corresponding to 80% yield. Intriguingly, we observed only trace formation of the monoacylated lipid product, including at early time points. Even when using substoichiometric amounts of palmitoyl-CoA, we primarily observed formation of the diacylated compound **4** with minimal formation of monoacylated cysteine (Figure S5). We attribute our observation to the rate-limiting step being the first acylation. Once acylated, the lipid likely forms mixed micelles with palmitoyl-CoA, positioning another thioester in proximity to the reactive thiol, leading to a more rapid second acylation. Such cooperative lipidation mechanisms may have played an important role in generating diacylated lipid species in the absence of membrane-bound enzymes during the early evolution of cellular life.

Microscopy studies were conducted to determine if dipalmitoylated-cysteine lipids could form membrane-bound vesicles. We observed small (<2 μm) lipid structures formed by self-assembly of **4** (Figure S6), which falls within a favorable protocell size range for supporting primitive biochemical processes (*27*). Vesicle formation was confirmed by cryo-electron microscopy (cryo-EM) (Figure 2d). The small size of the vesicles observed may be attributed to the fully saturated palmitoyl tails of lipid **4**, which will restrict membrane fluidity and influence vesicle assembly (*28*). Indeed, the phase transition temperature of membranes of **4** was found to be 60.4 °C (Figure S6). However, due to the limitations of optical microscopy in visualizing submicron vesicles, we sought to obtain larger vesicles to facilitate experimental observation and characterization. At low concentration, (10-15 mol% compared to the lipid), sterols such as cholesterol can disrupt the long-range lateral order of saturated lipids and help fluidize membranes (*29, 30*). Therefore, we hypothesized that addition of low concentrations of cholesterol would support the assembly of larger vesicles from lipid **4**. We found that addition of 10 mol% of cholesterol during the synthesis of **4** leads to the formation of vesicles between 1-5 µm in diameter that were easily observable by phase-contrast and fluorescence microscopy using 0.1 mol% BODIPY-FL (Figures 2e-f, and S6). As a control, we verified that palmitoyl-CoA, cysteine, cholesterol, or any combination of these components did not lead to membrane formation (Figure S7). Vesicles formed from **4** were able to encapsulate small polar molecules such as 8-hydroxypyrene-1,3,6-trisulfonic acid (HPTS), a highly polar fluorescent dye (Figure 2g). Similarly, synthesis of **4** in the presence of green fluorescent protein (GFP) also led to spontaneous entrapment of the protein, indicating that larger macromolecules can be encapsulated during vesicle formation (Figure 2h). The ability to encapsulate proteins during de novo lipid vesicle formation suggests that early protocells may have been able to capture primitive metabolic networks in a confined environment to sustain biochemical activity and couple metabolism with compartmentalization (*31, 32*).

Since our goal was to generate lipid membranes from simple metabolic precursors using a minimal set of soluble enzymes in the absence of preexisting membranes or lipid nanostructures, we next explored if the biosynthesis of palmitoyl-CoA **3** could be combined with chemical diacylation of cysteine for chemoenzymatic de novo membrane formation (Figure 3). Optimization of the reaction conditions allowed both the enzymatic and chemical reactions to occur simultaneously. Briefly, cysteine was added to a reaction mixture containing acetate, and the three enzymes ACS, ACC, FAS with the necessary cofactors, and incubated at 37 °C. The formation of lipid **4** was monitored by HPLC-ELSD-MS, reaching a yield of 43% after 12 hours. Small lipid structures (<2μm) were detected by phase-contrast microscopy (Figure S8), and cryo-EM analysis revealed the formation of nanometer sized bilayer vesicles (Figure 3b). Addition of 10 mol% cholesterol (relative to the final concentration of lipid **4**) to the reaction mixture led to the formation of larger vesicles with diameters reaching up to 5 μm (Figures 3c-d). As in the case of vesicles formed from chemically synthesized lipid **4** (Figure 2g-h), de novo vesicles generated during the chemoenzymatic reaction were able to spontaneously encapsulate HPTS (Figure S8) and GFP (Figure 3d) if these molecules were present in the buffer at the start of the reaction. The bilayer thickness for the de novo vesicles was estimated to be 2.4 nm based on cryo-EM analysis (Figure S9).

**Figure 3.**
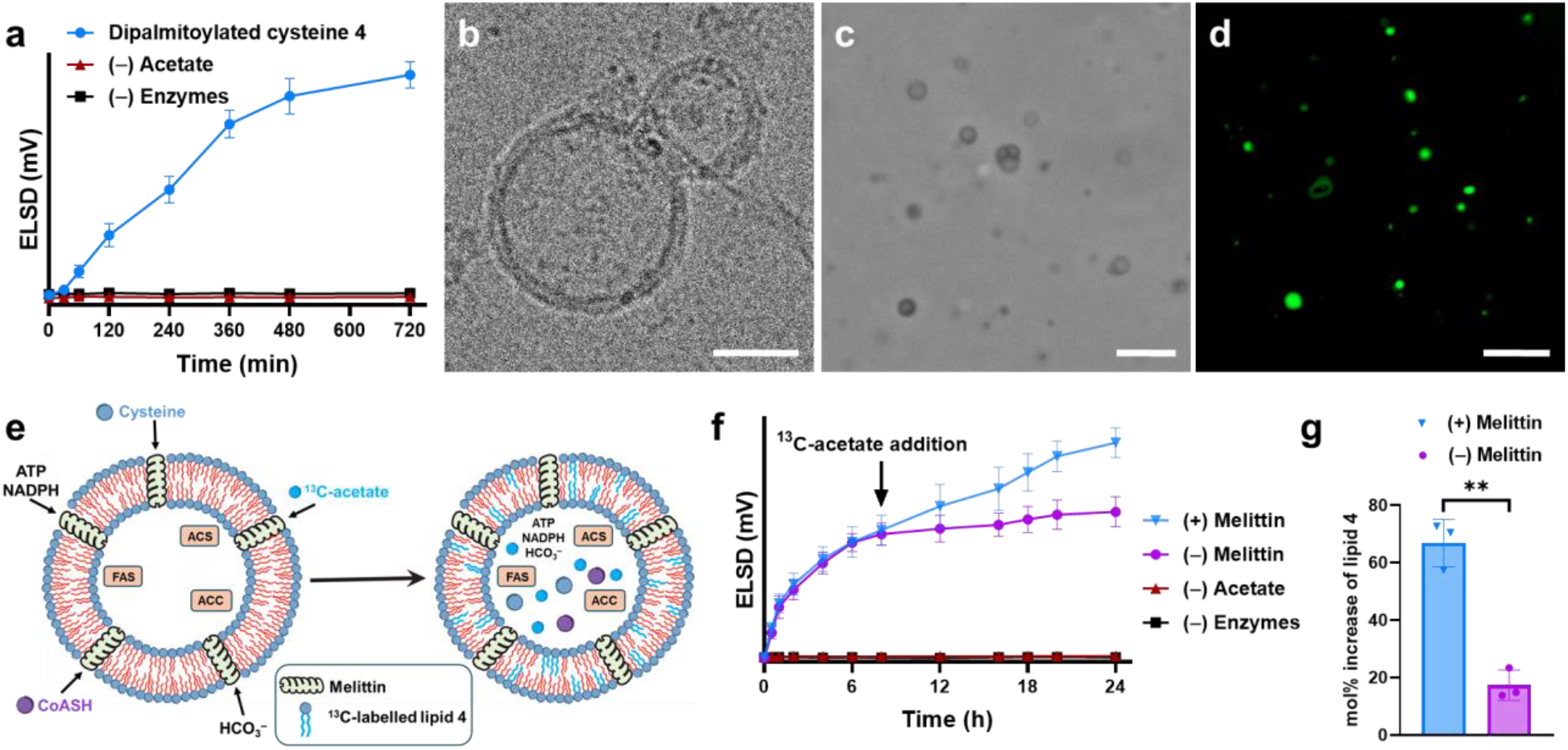
Chemoenzymatic synthesis of dipalmitoylated cysteine lipid 4 leading to de novo formation of vesicles. **(a)** Chemoenzymatic de novo generation of lipid **4** over 12 h. Negative controls show that no product is formed in the absence of acetate or without the enzymes ACS, ACC, and FAS. Data points presented as means ± s.d. (*n* = 3 technically independent samples). **(b)** Cryo-EM image of membrane-bound vesicles of lipid **4** generated de novo chemoenzymatically. Scale bar = 20 nm. **(c)** Phase-contrast microscopy image of membrane-bound vesicles of lipid **4** in the presence of 10 mol% cholesterol. Scale bar = 5 µm. **(d)** Spontaneous GFP encapsulation during the chemoenzymatic de novo formation of lipid **4** membranes in the presence of 10 mol% cholesterol. Scale bar = 5 µm. **(e)** Schematic representation of melittin-mediated pore formation in dipalmitoylated cysteine lipid **4** vesicles. New isotopically labeled lipid **4** (depicted in light blue) is synthesized after addition of fresh precursors that enter the vesicles via the melittin pores. **(f)** Chemoenzymatic de novo formation of lipid **4** monitored over time. 10 mol% melittin was added after 8 h of diacylation and incubated for 1 h. Fresh reaction precursors including ^13^C-acetate and cysteine were added and the reaction followed for a further 8 h. As shown in the graph, significant additional ^13^C-labeled lipid **4** is synthesized when melittin is present. Negative control without melittin addition showed reduced formation of lipid **4**. No lipid formation was observed in controls lacking acetate or enzymes. Data points presented as means ± s.d. (*n* = 3 technically independent samples). **(g)** Increase in the amount of lipid **4** after the addition of new reaction precursors in the presence and absence of melittin. Data points presented as means ± s.d. (*n* = 3 technically independent samples), and the significance determined using an unpaired t-test (two tailed). ***P = 0.0009.

Despite the presence of competing reactive species such as the acetyl-CoA and malonyl-CoA thioesters, cysteine underwent selective diacylation with palmitoyl-CoA to form **4**. No products bearing acetylated or malonylated tails were detected by HPLC-ELSD (Figure S10). One possible explanation is that the consumption of acetyl-CoA and malonyl-CoA by enzymes such as ACS, ACC, and FAS to produce palmitoyl-CoA occurs more rapidly compared to the much slower NCL-mediated diacylation reaction (Figure 2a, c). We also found that the acylation of cysteine with amphiphilic palmitoyl-CoA is strongly favored over acylation with more polar thioesters such as acetyl-CoA and malonyl-CoA (Figure S10). This preference may be attributed to the nature of cysteine (*33*), which could promote its partitioning into palmitoyl-CoA micelles, thus contributing to the observed chemoselective acylation.

We hypothesized that if the enzyme catalysts are entrapped within the de novo formed vesicles, membrane synthesis could be sustained internally. However, to enable continued lipid synthesis, there would also need to be mechanisms by which small molecule precursors can reach the interior of the protocells. Melittin is a short 26-amino acid soluble lytic peptide from bee venom that induces concentration dependent pore formation in lipid membranes (*34–37*). At low concentrations, melittin associates with the surface of lipid membranes without compromising their integrity, but at higher concentrations it inserts into the membrane and induces channel formation (*38*). In phospholipid membranes, the pore sizes formed by melittin are between 2.6-4.8 nm in diameter which lets small molecules, such as fluorescent dyes, to travel freely through the membranes while preventing the translocation of larger macromolecules such as enzymes (*39*). We sought to investigate whether melittin could similarly interact with de novo formed membranes composed of lipid **4**, generating pores that permit the influx of small molecules while retaining encapsulated enzymes within the vesicles. We prepared vesicles of **4** in reaction buffer containing HPTS and removed the excess unencapsulated dye via spin-filtration (Figure S11). We then tested dye leakage in response to increasing amounts of added melittin (0-25 mol%). Upon addition of 10 mol% melittin, vesicles lost their internal fluorescence within 1 hour of incubation, indicating membrane permeabilization, while retaining theirmorphology (Figure S11). In contrast, addition of 25 mol% melittin caused complete membrane disruption, as vesicles could no longer be visualized by microscopy, although lipid **4** was still present in the sample, as confirmed by HPLC-ELSD-MS analysis (Figure S12). As a control, in the absence of melittin, no leakage of HPTS was observed over a 16-hour period, supporting the role of melittin in membrane pore formation (Figure S13). Finally, analogous experiments with vesicles encapsulating GFP showed that the encapsulated protein was retained in the presence of 10 mol% melittin, indicating that melittin pores only allow the passage of small molecules across membranes while excluding larger macromolecules (Figure S13).

After demonstrating that short peptides can insert into the chemoenzymatically generated membranes and form nanochannels, we investigated if subsequent addition of reactive precursors would lead to additional lipid synthesis (Figure 3e). Since macromolecules like GFP are encapsulated during de novo protocell formation and show no significant leakage after melittin treatment, we reasoned that the enzymes ACS, ACC, and FAS are likewise encapsulated and retained inside the newly formed lipid **4** vesicles (Figure 3e). We chemoenzymatically formed **4** in the presence of HPTS, confirmed vesicle assembly by microscopy, and then removed non-encapsulated HPTS, enzymes, and other reactants by spin-filtration. We then added 10 mol% melittin to the vesicle population and incubated at 37 °C for 1 hour. Following confirmation of pore formation, indicated by HPTS leakage, we supplemented the system with fresh reagents and cofactors including cysteine, ^13^C-labeled acetate, CoASH, HCO_3_^−^, ATP, NADPH, and incubated for 8 hours. Lipid formation was monitored by HPLC-ELSD, revealing the formation of ^13^C-labeled lipids (Figure 3f), corresponding to a 67 mol% increase in total lipid content (Figure 3g). In contrast, in the absence of melittin, lipid levels increased by only 17 mol% over the same period (Figure 3g). The significant difference in lipid synthesis with and without melittin is likely due to the limited permeability of the vesicle membrane, as the newly added reaction precursors would have difficulty entering the vesicles simply through passive diffusion.

While cysteine offers a highly simple polar substrate for diacylation and membrane lipid formation, cellular membrane lipids are more complex, often bearing sugars or phosphorylated polar head groups. To better mimic the structural features of natural lipids, we considered alternative polar head groups with aminothiol reactive groups that might also be diacylated to form more complex lipid structures. An essential function of lipid membranes in cells is their maintenance of proton gradients, a requirement for chemiosmosis (*40*). However, vesicles formed from lipids containing acidic headgroups have difficulty supporting proton gradients due to proton shuttling across the membrane (Figure 4a). The need for proton gradient control suggests that more complex polar lipids developed early during the emergence of life, bridging prebiotic chemistry and early protocells (*40, 41*). We speculated that introducing a non-ionizable sugar head group rather than a carboxylic acid head group could render the resulting lipid membranes capable of maintaining proton gradients (Figure 4b).

**Figure 4.**
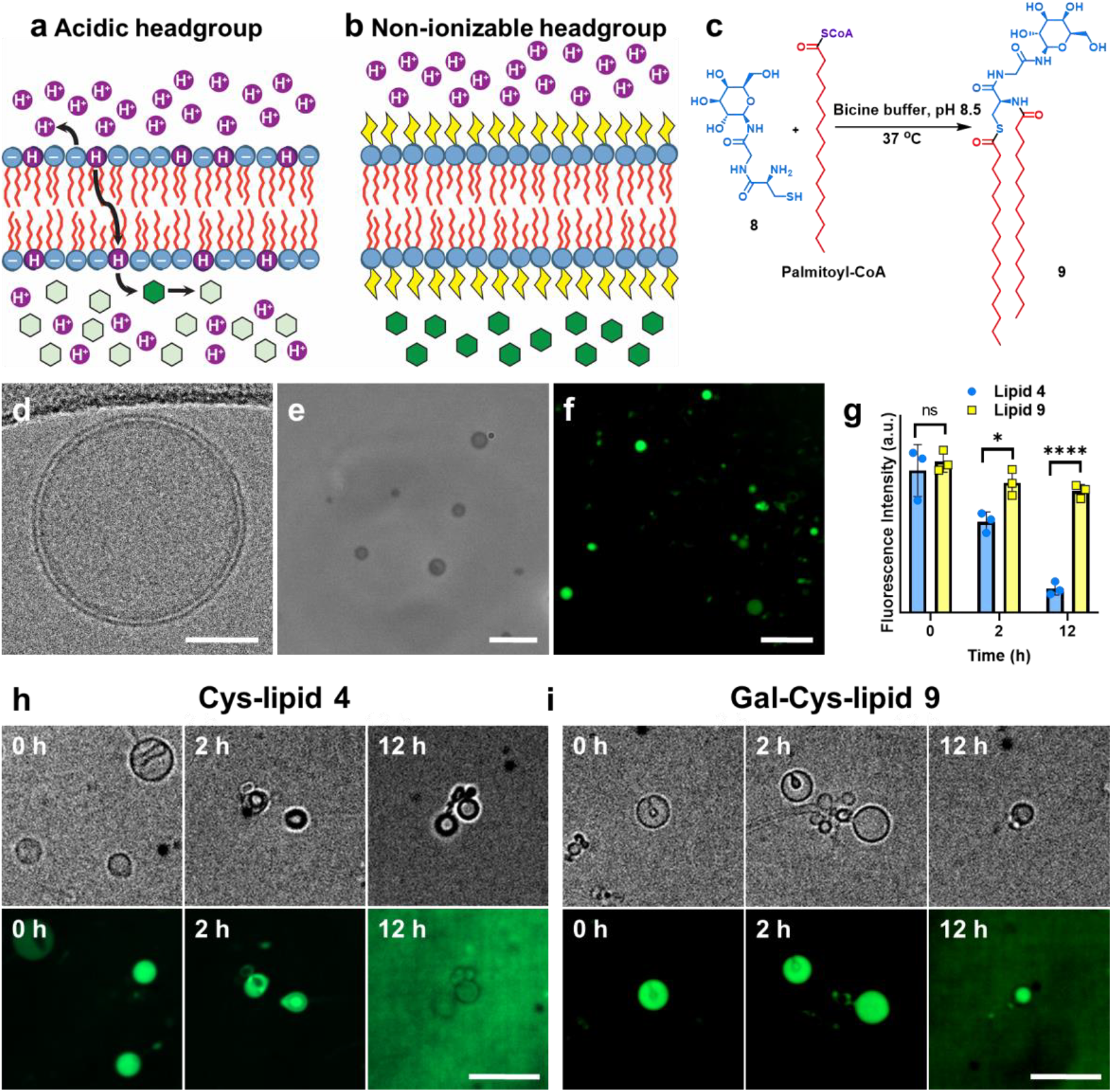
De novo formation of vesicles capable of maintaining a proton gradient. **(a-b)** Representative illustrations of a proton gradient across membranes containing acidic **(a)** versus non ionizable **(b)** lipid headgroups. Proton shuttling across the membrane is facilitated by protonation and flip-flop of the acidic lipid, while in membranes with non-ionizable headgroups, protons must cross the membrane in their charged form, which is energetically less favorable. **(c)** Reaction scheme for lipid **9** synthesis from palmitoyl-CoA and galacto-HG **8** in bicine buffer pH 8.5. **(d)** Cryo-EM image of a vesicle of lipid **9** generated from palmitoyl-CoA and **8** in the presence of 10 mol% cholesterol. Scale bar = 20 nm. **(e)** Phase-contrast microscopy image of de novo generated vesicles formed through the chemoenzymatic reaction using acetate and **8** as precursors, in the presence of 10 mol% cholesterol. Scale bar = 5 µm. **(f)** Fluorescence microscopy image of vesicles of **9** formed through the chemoenzymatic reaction showing encapsulation of GFP in the aqueous core. Scale bar = 5 µm. **(g)** Time-dependent decay of pH gradient in vesicles composed of either lipid **4** or **9**. Background-subtracted fluorescence intensity of encapsulated HPTS was measured at 0 h, 2 h and 12 h after buffer exchange from bicine (pH = 8.5) to citrate (pH = 4.5). Data points are presented as means ± s.d. (*n* = 3 technically independent samples), and the significance determined using an unpaired t-test (two tailed). 0 h, P = 0.6130; 2 h *P = 0.0143; 12 h ****P < 0.0001. **(h-i)** Brightfield (*top*) and fluorescence (*bottom*) microscopy images showing pH gradient decay in vesicles composed of lipid **4 (h)** and galactolipid **9 (i)** at 0 h, 2 h, and 12 h after buffer exchange. Scale bars = 5 µm.

To test this, we synthesized a short glycopeptide polar head group galactosaminothiol (galacto-HG) **8** (Scheme S1) and showed that spontaneous diacylation and vesicle formation occurs in the presence of palmitoyl-CoA to form non-canonical galactolipid **9** (Scheme S2, Figures 4c, and S14). Galacto-HG **8** undergoes diacylation during the enzymatic synthesis of palmitoyl-CoA to generate galactolipid **9**. Optimization of the reaction conditions enabled concurrent synthesis of enzymatically generated palmitoyl-CoA, diacylation of galacto-head group **8**, and vesicle formation (Figures 4d-e). Vesicles formed from galactolipid **9** were shown to be able to encapsulate small molecules (Figure S15), and proteins (Figure 4f), and they could support additional lipid synthesis in the presence of pore forming peptide melittin, similar to vesicles formed from **4** (Figure S16).

To assess whether membranes formed from either lipid **4** or lipid **9** differ in their ability to maintain proton gradients, we generated vesicles de novo using cysteine or galacto-HG **8** as reactive head groups in the presence of HPTS in bicine buffer (pH 8.5) and subsequently triggered a proton gradient by exchanging the external buffer for citrate buffer (pH 4.6). The HPTS dye exhibits high fluorescence in alkaline media, but its fluorescence is quenched by more than 99% in more acidic environments (*42*). Following buffer exchange, fluorescence measurements indicated that a proton gradient was initially maintained across both types of membranes. However, the fluorescence intensity of the vesicles decreased over time in both cases, likely reflecting acidification of the internal lumen of the vesicles due to proton influx (Figure 4g). In vesicles composed of dipalmitoylated-cysteine **4**, fluorescence decreased by 83±6% over 12 hours, whereas vesicles formed by galactolipid **9** showed only a 19±7% decrease over the same time period (Figures 4g). Confocal microscopy images corroborated the fluorescence quenching observed for lipid **4** vesicles (Figure 4h), while confirming the persistence of fluorescence in lipid **9** vesicles even after 12 hours (Figure 4i). Lipid **4** possesses a small, acidic polar head group, which may facilitate the shuttling of protons from the outer buffer to the inner lumen of the membranes after buffer exchange, similar to observations with single-chain fatty acids, albeit occurring at a slower rate due to the presence of two saturated and long-chain fatty-acyl hydrocarbon tails (*43, 44*). Indeed, control experiments performed using oleic acid vesicles showed a much more rapid drop in fluorescence upon buffer exchange (Figure S17). In contrast, galactolipid **9** contains a galactosamino moiety as the head group, which does not possess an easily ionizable functional group and is significantly larger in size. The ability to maintain and harness proton gradients could have conferred protocells with competitive advantages and driven the evolution of more complex membrane systems.

## Conclusion

In summary, we demonstrate that de novo vesicle formation can take place in the absence of preexisting membranes or lipid templates, upending a long-standing dogma in cellular biology. Non-canonical diacylated cysteine membranes of **4** can form from soluble enzymes and are fueled by primitive non-lipidic metabolites such as acetate, and cysteine. We show that palmitoyl-CoA enzymatically generated from acetate can spontaneously and chemoselectively couple with cysteine to generate lipid dipalmitoylated-cysteine **4**. The resulting membrane lipids spontaneously self-assemble into vesicles. Simple peptides can form stable pores in de novo formed membranes, allowing for efficient shuttling of fresh acetate and cysteine to generate additional membrane lipids in the system. We could adapt this general strategy to generate a more complex biomimetic galactolipid **9** which forms membranes de novo that maintain proton gradients. Our study reveals how soluble enzymes and primitive metabolites can drive the formation of lipid membranes, which are relatively complex biophysical structures. A conceptually similar process may have served as an intermediate step in the evolution of life, linking simplified prebiotic biochemistry to the first living cells. While extant cell membranes possess lipids with more complex head groups, it is possible that earlier protocell membrane head groups were composed of simpler functional groups that were later elaborated to improve membrane properties, such as decreasing permeability to small molecules and ions, which would have been necessary for effective chemiosmosis. Beyond implications for the origin of life, our work also suggests new routes for encoding for membrane formation in artificial cells and new ways to generate lipid structures de novo for applications in nanotechnology, medicine, and materials science. We anticipate that exploring alternative enzymes as well as reactive head groups could lead to a variety of lipids that assemble to form membranes with diverse physical properties.

## Acknowledgments

This material is based upon work supported by the National Science Foundation (Award MCB-2124105). The authors acknowledge the facilities along with the scientific and technical assistance of the staff of the cryo-EM facility at UC San Diego.

