## Supplementary Information for "De Novo Protocell Membrane Formation Fueled by Primitive Metabolites"

**Table of Contents**

1. **General Methods, Instrument Details and Materials**
2. **Experimental Procedures**
3. **Supplementary Schemes**
4. **Supplementary Figures**
5. **Characterization Spectra**
6. **References**

### 1. General Methods, Instrument Details and Materials

*N*-Boc-*L*-Cys(Trt)-COOH, Borane-tetrahydrofuran ($BH_3$·THF), anhydrous tetrahydrofuran (THF), methanol (MeOH), *N*,*N*-diisopropylethylamine (DIPEA), anhydrous chloroform ($CHCl_3$), pyridine ($C_5H_5N$), trifluoroacetic acid (TFA), acetic acid ($CH_3COOH$), and tris(2-carboxyethyl)phosphine hydrochloride (TCEP) were obtained from Sigma-Aldrich. Texas Red® 1,2-dihexadecanoyl-*sn*-glycero-3-phosphoethanolamine, triethylammonium salt (Texas Red® DHPE) was obtained from Life Technologies. BODIPY FL DHPE was obtained from ThermoFisher Scientific. Deuterated chloroform ($CDCl_3$) and methanol ($CD_3OD$) were obtained from Cambridge Isotope Laboratories. Commercially available standards of acetyl coenzyme A (Acetyl-CoA), malonyl coenzyme A (Malonyl-CoA), as well as nicotinamide adenine dinucleotide phosphate (NADPH) was purchased from Sigma-Aldrich. Acetyl-CoA carboxylase human (ACC) enzyme was purchased from Sigma-Aldrich (#A6861). AMP-Glo™ assay was acquired from Promega. Acetyl-Coenzyme A assay kit was ordered from Sigma-Aldrich. ELISA assay kit for malonyl-CoA was acquired from MyBioSource. All reagents obtained from commercial suppliers were used without further purification unless otherwise noted.

Solvent mixtures for chromatography are reported as v/v ratios. HPLC analysis was carried out on an Eclipse Plus C8 analytical column with *Phase A*/*Phase B* gradients [*Phase A*: $H_2O$ with 0.1% formic acid; *Phase B*: MeOH with 0.1% formic acid]. The HPLC was equipped with a UV-vis detector, a 380 Varian-Agilent evaporative light scattering detector (ELSD), and a 6120 Agilent Quadrupole MS (MS). HPLC purification was carried out on Zorbax Eclipse Plus C8 preparative column with *Phase A*/*Phase B* gradients [*Phase A*: $H_2O$ with 0.1% formic acid; *Phase B*: MeOH with 0.1% formic acid].

Proton nuclear magnetic resonance ($^1H$ NMR) spectra were recorded on a Jeol ECA-500 (Jeol Ltd., Tokyo, Japan) or a Jeol ECA-400MHz spectrometer and were referenced relative to residual proton resonances in $CDCl_3$ (at δ 7.24 ppm) or $CD_3OD$ (at δ 4.87 or 3.31 ppm). Chemical shifts were reported in parts per million (ppm, δ) relative to tetramethylsilane (δ 0.00). $^1H$ NMR splitting patterns are assigned as singlet (s), doublet (d), triplet (t), quartet (q) or pentuplet (p). All first-order splitting patterns were designated on the basis of the appearance of the multiplet. Splitting patterns that could not be readily interpreted are designated as multiplet (m) or broad (br). Carbon nuclear magnetic resonance ($^{13}C$ NMR) spectra were recorded on a Jeol ECA-400MHz spectrometer, and were referenced relative to residual proton resonances in $CDCl_3$ (at δ 77.23 ppm) or $CD_3OD$ (at δ 49.15 ppm). Electrospray Ionization-Time of Flight (ESI-TOF) spectra were obtained on an Agilent 6230 Accurate-Mass TOF-MS mass spectrometer.

Light microscopy images were acquired using an Olympus BX51 optical microscope with a 100x objective, with bright field imaging conducted in phase-contrast mode. Spinning-disk confocal microscopy images were acquired on a Yokagawa spinning-disk system (Yokagawa, Japan) built around an Axio Observer Z1motorized inverted microscope (Carl Zeiss Microscopy GmbH, Germany) with a 63x, 1.40 NA oil immersion objective to an Evolve 512x512 EMCCD camera (Photometrics, Canada) using ZEN imaging software (Carl Zeiss Microscopy GmbH, Germany). Bright field imaging on the confocal system was performed in differential interference contrast (DIC) mode. The fluorophores were excited with a 20 mW DPSS laser (Texas Red®).

Cryo-electron microscopy grids (Lacey Carbon, Electron Microscopy Sciences #LC300-Cu) were glow-discharged using a Emitech K350 unit at 20 mA for 30 s. A Vitrobot (Mark IV, Thermo Fisher Scientific) was used to plunge the cryo grids into liquid ethane (3.0 µL sample per grid). For the vesicle formation starting from palmitoyl-CoA, images were collected on a Titan Krios G3 (Thermo Fisher Scientific) operated at 300 keV equipped with a K3 detector and a 1067HD BioContinuum energy filter (Gatan) with a 15 eV slit width. Images were acquired with a total dose of 50 e/Å2 at 2.16 Å/pixel and 4 µm nominal defocus. For the vesicle formation reactions with melittin incubation, images were acquired on a Talos Arctica (FEI) operated at 200 kV and collected with a total dose of 40 e/Å2 at 0.95 Å/pixel and a 3 µm nominal defocus.

NanoDrop 2000C spectrophotometer was used for UV-vis measurements. Fluorescence, luminescence and absorbance measurements were carried out on a Tecan infinite F200 plate reader instrument. Phase transition temperature experiments were conducted on a Nano DSC from TA instruments, Waters.

### 2. Experimental Procedures

Choice of enzymes for the synthesis of palmitoyl-CoA

To enable the synthesis of palmitoyl-CoA, a set of three enzymes was selected. First, ACS from *E. coli* was selected as it is well-studied and reported to be efficient (*1*). ACS activates acetate with CoA, consuming ATP and releasing AMP to produce acetyl-CoA. Second, the commercially available human acetyl-CoA carboxylase 2 (ACC) was chosen for generating malonyl-CoA from acetyl-CoA, also in an ATP-dependent process. Finally, the type I fatty acid synthase from *Corynebacterium glutamicum* (FAS) was selected due to its reported ability to efficiently generate palmitoyl-CoA from acetyl-CoA and malonyl-CoA.

Expression and purification of ACS

The plasmid encoding for $His_6$-tagged wild-type *E. coli* ACS (pETM11) was purchased from Addgene (*1*). Competent *E. coli* BL21(DE3) cells were transformed with the pETM11 plasmid and grown overnight at 37 °C in Luria-Bertani (LB) broth containing 0.1 mg/mL of kanamycin. Afterwards, 1 mL of the overnight culture was used to inoculate 1 L of freshly autoclaved LB medium containing 0.1 mg/mL of kanamycin. The rest of the overnight culture was stored as 25% glycerol stocks at –80 °C. The culture was grown at 37 °C in a shaker-incubator until the $OD_{600}$ reached 0.6. Overexpression of ACS was induced by addition of 1 mM isopropyl 1-thio-D-galactopyranoside (IPTG). The cells were then grown for 16 h at 18 °C, after which they were harvested by centrifuging at 6000 rcf for 20 min at 4 °C. The pellet was resuspended by vortexing in 10 mL of lysis buffer containing 25 mM Tris buffer, pH 8.0, 0.5 M NaCl, 2 mM β-mercaptoethanol, 1 mg/mL lysozyme and a cocktail of protease inhibitors (SigmaFast®). Following cell lysis by an ultrasonicator probe, debris was removed by centrifuging (13000 rcf, 20 min, 4 °C). The supernatant was incubated in a gravity column with $Ni^{2+}$-nitrilotriacetate (Ni-NTA) HisPur agarose resin pre-equilibrated with 10 mM imidazole on a shaker table for 3 h at 4 °C. Next, the flow-through was discarded and the resin was washed with 10 mL each of 20 mM imidazole and 50 mM imidazole solutions. Finally, $His_6$-tagged ACS was eluted with 250 mM imidazole and collected in 1 mL fractions. The fractions were analyzed by SDS-PAGE to check for impurities, and subsequently concentrated in a centrifugal filter with a 10000 nominal molecular weight limit (Amicon Ultra-4, Merck Millipore) after buffer exchange to remove excess imidazole. The sample was further purified by size-exclusion chromatography (column: Superose 6 10/300GL, GE Healthcare, buffer G: 100 mM $Na_2HPO_4/NaH_2PO_4$ pH 7.4, 100 mM NaCl) and examined for its oligomeric state. Fractions were pooled, concentrated and aliquoted to a final concentration of 0.87 µM of protein containing 10% glycerol each and stored at –80 °C. Protein $A_{280}$ and concentration measurements were carried out using a NanoDrop (ThermoFisher Scientific).

Expression and purification of FAS

Expression and purification protocol of FAS was performed as previously described (*2*). In short, competent *E. coli* BL21 cells were transformed with the pBbE5c plasmid and grown overnight at 37 °C in LB broth containing 0.1 mg/mL of chloramphenicol. Afterwards, 1 mL of the overnight culture was used to inoculate 1 L of freshly autoclaved LB medium containing 0.1 mg/mL of chloramphenicol. The rest of the overnight culture was stored as 25% glycerol stocks at –80 °C. The culture was grown at 37 °C in a shaker-incubator till the $OD_{600}$ reached 0.65. Overexpression of FAS was induced by addition of 1 mM isopropyl 1-thio-D-galactopyranoside (IPTG). The cells were then grown for 16 h at 18 °C, after which they were harvested by centrifuging at 6000 rcf for 20 min at 4 °C. The pellet was resuspended by vortexing in 10 mL of lysis buffer containing 25 mM Tris buffer, pH 8.0, 0.5 M NaCl, 2 mM β-mercaptoethanol, 1 mg/mL lysozyme and a cocktail of protease inhibitors (SigmaFast®). Following cell lysis by an ultrasonicator probe, debris was removed by centrifuging (13000 rcf, 20 min, 4 °C). The supernatant was incubated in a gravity column with

$Ni^{2+}$-nitrilotriacetate (Ni-NTA) agarose resin pre-equilibrated with 10 mM imidazole on a shaker table for 2 h at 4 °C. Next, the flow-through was discarded and the resin was washed with 10 mL each of 20 mM imidazole and 50 mM imidazole solutions. Finally, $His_6$-tagged FAS was eluted with 250 mM imidazole and collected in 1 mL fractions. The fractions were analyzed by SDS-PAGE to check for impurities, and subsequently concentrated in a centrifugal filter with a 100000 nominal molecular weight limit (Amicon Ultra-4, Merck Millipore). The sample was further purified by size-exclusion chromatography (column: Superose 6 10/300GL, GE Healthcare, buffer G: 100 mM $Na_2HPO_4/NaH_2PO_4$ pH 7.4, 100 mM NaCl) and examined for its oligomeric state. Fractions were pooled, concentrated and aliquoted to a final concentration of 1.83 µM of protein containing 10% glycerol each and stored at –80 °C. Protein $A_{280}$ and concentration measurements were carried out using a NanoDrop (ThermoFisher Scientific).

Enzymes activity assays

Enzyme activity was assessed using both indirect (reagents consumption) and direct (product formation) methods. Indirect measurements for ACS and ACC were based on the quantification of the AMP formed using the AMP-Glo™ assay kit (Promega), while FAS activity was evaluated by monitoring NADPH consumption. Direct quantification of products was performed using an acetyl-CoA assay kit (Sigma Aldrich) for ACS activity, an ELISA-based assay to quantify the malonyl-CoA formed in the presence of ACC, and HPLC-ELSD for palmitoyl-CoA generated in the FAS-catalyzed reaction.

AMP-Glo™ activity assay

The AMP-Glo™ assay is a bioluminescent method to determine the concentration of AMP in a biochemical reaction. The assay contains two reagents: the first removes any remaining ATP, and converts the AMP produced into ADP; the second reagent (AMP Detection Solution) drives the conversion of ADP to ATP, and the detection of ATP through a luciferase reaction. The amount of AMP produced by the reaction is proportional to the light measured and can be extrapolated using a standard curve. Assay standards and buffers were prepared according to the protocol provided in the kit.

Generating an AMP Standard Curve

AMP standards were prepared in a 384-well plate in triplicate. A 2000 µM AMP solution was used as the starting concentration, and six additional standards were prepared by twofold serial dilution across the plate, resulting in final concentrations of 2000, 1000, 500, 250, 125, 62.5, and 31.25 µM. One well containing only dilution buffer served as the negative control. After adding the AMP-Glo™ reagents according to the manufacturer's instructions, luminescence was measured using a Spark multimode plate reader (Tecan), and the signal intensity was plotted against AMP concentration to generate a standard curve. The difference in relative luminescence units (DRLU) between the luminescence observed at each concentration and the in the blank solution (with no-AMP) was calculated. The DRLU values were plotted on the Y-axis against their corresponding AMP concentrations on the X-axis to generate the standard curve (Figure S1a).

AMP-Glo™ assay for ACS activity

Acetyl-CoA is generated from acetate and coenzyme A in an ATP-dependent reaction that produces AMP as byproduct. To test the activity of ACS, the AMP detection assay was carried out on a 50 µL scale. First, acetyl-CoA formation reaction was set up in a glass vial in bicine buffer at pH 8.5. The reaction mixture contained 30 µL of 100 mM bicine buffer (containing 10 mM TCEP), 5 µL of 10 mM acetate (in sterile $H_2O$), 5 µL of 10 mM coenzyme A (CoASH) (in sterile $H_2O$), 5 µL of 10 mM ATP (in sterile $H_2O$), and 5 µL of ACS (10 µM in elution buffer) at 37 °C. After 1 h of reaction, 5 µL of the reaction mixture was added to a 384-well plate along with 20 µL of AMP detection solution, 10 µL of AMP-Glo Reagent 1 (AMP-Glo 1), and 5 µL of 2X substrate (also provided with the kit). The solutions were prepared as suggested in the protocol provided with the kit. After adding AMP-Glo 1 to all wells, the plate was mixed by shaking for 1-2 minutes. The plate was then incubated at room temperature for 1 h. After that, 20 µL of AMP detection solution was added to all wells. The plate was then incubated at rt for 1 h after mixing. The luminescence was measured using a Spark multimode plate reader (Tecan). The concentration of AMP was extrapolated using the standard curve described above (Figure S1a). The AMP concentration determined in the assay corresponded to 650±40 µM. The reactions were conducted in triplicates (Figure S1c). Two control conditions were prepared following the same protocol, with the omission of either ACS or acetate to confirm that AMP production was dependent on both enzyme and substrate (Figure S1c).

Acetyl-CoA detection assay

For direct assessment of ACS enzymatic activity, acetyl-CoA production was measured using an acetyl-CoA assay kit (Sigma Aldrich). This kit employs a coupled enzymatic reaction that produces a fluorometric signal ($\lambda_{ex}$ = 535 nm, $\lambda_{em}$ = 587 nm), which is proportional to the amount of acetyl-CoA generated. Standards and buffers were prepared following the manufacturer's protocol.

Generating an acetyl-CoA standard curve

Acetyl-CoA standards were prepared in triplicate. A 2 mM standard solution of acetyl-CoA was used to generate final concentrations of 0 (blank), 200, 400, 600, 800, and 1000 µM standards. The standards were treated with the acetyl-CoA assay kit described above, fluorescence was recorded and the values were plotted on the Y-axis against their corresponding acetyl-CoA concentrations on the X-axis to generate the standard curve (Figure S1b).

Acetyl-CoA detection assay for ACS activity

Acetyl-CoA forming reactions were set up in glass vials in bicine buffer at pH 8.5. Each reaction was set up in 20 µL of 100 mM bicine buffer (containing 10 mM TCEP) with the addition of 5 µL of 10 mM acetate (in sterile $H_2O$), 5 µL of 10 mM coenzyme A (CoASH) (in sterile $H_2O$), 5 µL of 10 mM ATP (in sterile $H_2O$), and 5 µL of ACS (10 µM in elution buffer) at 37 °C. 10 µL of the reaction mixture weas added in triplicate to a 96 well plate. Samples were brought to a final volume of 50 µL with acetyl-CoA assay buffer. A blank was made for each sample as control by omitting the Reaction Initiator Solution (provided in kit) in the reaction mixture. To correct for background created by free coenzyme A and succinyl-CoA, 10 µL of acetyl-CoA quencher solution (provided in kit) was added to each sample, and sample blank. The reactions were incubated at room temperature for 5 min. Subsequently, 2 µL of quencher remover (provided in kit) was added, the samples were thoroughly mixed, and incubated for additional 5 min. Following preparation according to the assay protocol, the plate was protected from light using aluminum foil and incubated for 10 min at 37 °C. Finally, the fluorescence intensity was measured ($\lambda_{ex}$ = 535 / $\lambda_{em}$ = 587 nm). The concentration of acetyl-CoA was extrapolated using the standard curve described above. The concentration of acetyl-CoA determined using this assay corresponded to 660±25 µM (Figure S1d). The reactions were conducted in triplicate. Two control experiments were prepared following the same protocol, with the omission of either ACS or acetate to confirm that acetyl-CoA production was dependent on both enzyme and substrate.

AMP-Glo™ assay for ACC activity

To confirm the activity of the commercially acquired ACC, AMP formation was determined using the same method described for ACS. The malonyl-CoA synthesis reaction was carried out in a glass vial in bicine buffer at pH 8.5. Instead of acetate, 5 µL of 10 mM acetyl-CoA (in sterile $H_2O$) was used as the starting substrate, and 5 µL of ACC (10 µM in sterile $H_2O$) was used in place of ACS. The concentration of AMP was extrapolated from the AMP standard curve described above (Figure S1a). The AMP concentration determined with this assay corresponded to 756±20 µM (Figure S2a). The reaction was conducted in triplicate. Two control experiments were prepared following the same protocol, with the omission of either ACC or acetyl-CoA to confirm that malonyl-CoA production was dependent on both enzyme and substrate.

Malonyl-CoA detection assay

For direct assessment of ACC enzymatic activity, malonyl-CoA production was measured using a malonyl-CoA ELISA assay kit (MyBioSource). This assay uses the quantitative sandwich enzyme immunoassay method. Antibody specific for malonyl-CoA was pre-coated onto the microplate(s) provided in the kit. Standards and buffers were prepared following the manufacturer's protocol.

Generating a malonyl-CoA standard curve

Malonyl-CoA standards were prepared in triplicate. A 2 mM standard solution of malonyl-CoA was used to generate final concentrations of 0 (blank), 200, 400, 600, 800, and 1000 µM standards. The standards were analyzed by malonyl-CoA ELISA assay kit (MyBioSource) and the values obtained were plotted on the Y-axis against their corresponding malonyl-CoA concentrations on the X-axis to generate the standard curve (Figure S2b).

Malonyl-CoA ELISA detection assay for ACC activity

The malonyl-CoA formation reaction was carried out in a glass vial in bicine buffer at pH 8.5. The reaction mixture contained 35 µL of 100 mM bicine buffer (containing 10 mM TCEP), 5 µL of 10 mM acetyl-CoA (in sterile $H_2O$), 5 µL of 10 mM $HCO_3^-$ (in sterile $H_2O$), and 5 µL of ACC (10 µM in sterile $H_2O$) at 37 °C. After 1 h of reaction, 5 µL of the reaction mixture was added to the microplate wells provided with the kit. Samples were prepared according to the manufacturer's instruction and added to the wells, where any malonyl-CoA present was captured by the immobilized antibody. After removing any unbound substances, biotin-conjugated antibodies specific for malonyl-CoA was added to the wells, forming the "ELISA sandwich". After washing to remove unbound biotin-conjugated antibodies, avidin conjugated Horseradish Peroxidase (HRP) was added to the wells. Following a wash to remove any unbound avidin-HRP reagent, a tetramethylbenzidine (TMB) substrate solution was added to the wells and color developed in proportion to the amount of malonyl-CoA bound in the initial step. The color development was stopped using the Stop solution provided in the kit, and the intensity of the color was measured using a Spark multimode plate reader (Figure S2c). The malonyl-CoA concentration determined with the assay corresponded to 710±20 µM. The reaction was conducted in triplicate. Control experiments were carried out following the same procedure described above, but in absence of ACS and ACC, or acetate, respectively.

Malonyl-CoA ELISA detection assay for ACS and ACC activity combined

The malonyl-CoA formation reaction was carried out in a glass vial in bicine buffer at pH 8.5. The reaction mixture contained 20 µL of 100 mM bicine buffer (containing 10 mM TCEP), 5 µL of 10 mM acetate (in sterile $H_2O$), 5 µL of 10 mM CoASH (in sterile $H_2O$), 5 µL of 10 mM $HCO_3^-$ (in sterile $H_2O$), 5 µL of ACS (10 µM in sterile $H_2O$), 5 µL of ACC (10 µM in sterile $H_2O$) at 37 °C. After 1 h of reaction, 5 µL of the reaction mixture was added to the microplate wells provided with the kit. Samples were prepared according to the manufacturer's instruction and added to the wells, where any malonyl-CoA present was captured by the immobilized antibody. After removing any unbound substances, biotin-conjugated antibodies specific for malonyl-CoA was added to the wells, forming the "ELISA sandwich". After washing to remove unbound biotin-conjugated antibodies, avidin conjugated Horseradish Peroxidase (HRP) was added to the wells. Following a wash to remove any unbound avidin-HRP reagent, a tetramethylbenzidine (TMB) substrate solution was added to the wells and color developed in proportion to the amount of malonyl-CoA bound in the initial step. The color development was stopped using the Stop solution provided in the kit, and the intensity of the color was measured using a Spark multimode plate reader (Figure S2d). The malonyl-CoA concentration determined with the assay corresponded to 695±30 µM. The reaction was conducted in triplicate. Control experiments were carried out following the same procedure described above, but in absence of ACS and ACC, or acetate, respectively.

NADPH detection assay and standard curve

To test the activity of FAS, an NADPH consumption assay was carried out measuring the fluorescence emission at 470 nm (excitation 340 nm). NADPH standards were prepared in triplicate. A 2 mM standard solution of NADPH was used to generate final concentrations of 0 (blank), 100, 200, 300, 400, 500, 600, 700, 800, 900, 1000, 1100 µM standards. The standards were analyzed by fluorescence spectroscopy and the values obtained were plotted on the Y-axis against their corresponding NADPH concentrations on the X-axis to generate the standard curve (Figure S3a). The data was corrected for the NADPH autooxidation over time.

NADPH detection assay for FAS activity (*2*)

To test the activity of FAS, an NADPH consumption assay was carried out on a 50 µL scale. 25 µL of 100 mM bicine buffer, pH 8.5 (containing 10 mM TCEP), 10 µL of 10 mM acetyl-CoA (in sterile $H_2O$), 10 µL of 100 mM NADPH (in sterile $H_2O$) and 5 µL of FAS (50 µM in elution buffer) were tumbled at 37 °C. After 1 min of recording the emission at 470 nm using a Spark multimode plate reader (Tecan), the reaction was started by the addition of 10 µL of 7 mM malonyl-CoA and the emission was continuously measured for 1 h. The concentration of NADPH consumed in the reaction was extrapolated to be 1.20±52 mM, which corresponds to 570±20 µM palmitoyl-CoA being generated during the reaction (Figure S3b).

Generating a palmitoyl-CoA standard curve for HPLC-ELSD-MS quantification

Palmitoyl-CoA standards were prepared in triplicate. A 2 mM standard solution of palmitoyl-CoA was used to generate final concentrations of 0 (blank), 100, 200, 300, 400, 500, 600, 700, 800, 900, and 1000 µM standards. The standards were analyzed by HPLC-ELSD-MS and the values obtained were plotted on the Y-axis against their corresponding palmitoyl-CoA concentrations on the X-axis to generate the standard curve (Figure S3c). Chromatographic separation was performed with a mobile-phase system with gradients based on *Phase A* (water, 10 mM triethylamine/acidic acid buffer, adjusted to pH 9.0) and *Phase B* (acetonitrile). A multistep gradient was applied at a flow rate of 0.25 mL/min, starting with 7% *Phase B*, and linearly increasing it to 60% at 6 min, to 70% at 9.5 min and finally to 90% at 10 min runtime.

Palmitoyl-CoA quantification for FAS activity

Palmitoyl-CoA formation was quantified using HPLC-ELSD-MS analysis. The reaction was carried out on a 50 µL scale. 25 µL of 100 mM bicine buffer, pH 8.5 (containing 10 mM TCEP), 10 µL of 10 mM acetyl-CoA (in sterile $H_2O$), 10 µL of 100 mM NADPH (in sterile $H_2O$) and 5 µL of FAS (50 µM in elution buffer) were tumbled at 37 °C. The reaction was started by the addition of 10 µL of 70 mM malonyl-CoA. The reactions were conducted in triplicate. The concentration of palmitoyl-CoA was extrapolated from the standard curve described above. The palmitoyl-CoA concentration determined with this assay corresponded to 630±35 µM (Figure S3d). Two control experiments were prepared following the same protocol, with the omission of either FAS or acetyl-CoA to confirm that palmitoyl-CoA production was dependent on both enzyme and substrates.

Palmitoyl-CoA formation starting from acetate as carbon feedstock

Palmitoyl-CoA was generated starting from acetate using ACS, ACC, and FAS in a reaction cascade. Briefly, 14 µL of 1 M bicine buffer, pH 8.5 (containing 100 mM TCEP), 2.5 µL of 200 mM acetate (in sterile $H_2O$), 1.75 µL of 200 mM CoASH (in sterile $H_2O$), 1.75 µL of 200 mM $HCO_3^-$ (in sterile $H_2O$), 3.75 µL of 200 mM ATP (in sterile $H_2O$), 3.75 µL of 200 mM NADPH (in sterile $H_2O$), 5 µL of ACS (50 µM in elution buffer), 5 µL of 50 µM ACC (in sterile $H_2O$), and 12.5 µL of FAS (20 µM in elution buffer) were added to a glass vial and tumbled at 37 °C. The reaction was monitored for 4 h using HPLC-ELSD-MS (Figure S4). For each HPLC-ELSD-MS run, a small aliquot (10 µL) of the reaction was taken and centrifuged to remove protein debris before loading onto the column. The amount of palmitoyl-CoA generated was quantified using the standard curve described above and corresponded to 530±20 µM.

Synthesis of the Dipalmitoylated Cysteine **4**

To a stirred solution of cysteine (24 mg, 0.2 mmol, 1.00 equiv) and NaOH (16 mg, 0.4 mmol, 2.00 equiv) in water (100 µL), THF (400 µL) was added, then the reaction mixture was stirred at 0 ºC. Palmitoyl chloride (148 µL, 0.44 mmol 2.2 equiv) was added dropwise, then the reaction mixture was further stirred for 24 h. After the reaction, THF was removed under reduced pressure, then the mixture was acidified by addition of aqueous 6 M HCl solution. The mixture was extracted with EtOAc 3 times, dried over $MgSO_4$, filtered, then concentrated under reduced pressure. Dipalmitoylated cysteine **4** was obtained (29.83 mg, 35.2%) as a white powder after purification by flash column chromatography ($SiO_2$, hexane/EtOAc 90:10 to hexane/EtOAc. The product fractions were concentrated and analyzed by $^{1}H$ and $^{13}C$ NMR as well as HR-MS. $^{1}H$ NMR (400 MHz, $CDCl_3$): δ 6.69 (s, 1H), 4.69 (s, 1H), 3.36 (d, 2H), 2.62-2.58 (t, 2H), 2.26-2.22 (t, 2H), 1.63 (m, 4H), 1.25 (m, 48H), 0.89-0.86 (m, 6H). $^{13}C$ NMR (125 MHz, CDCl3): δ 201.82, 175.84, 171.90, 54.05, 44.11, 36.41, 32.07, 29.85, 29.82, 29.79, 29.75, 29.66, 29.63, 29.58, 29.52, 29.45, 29.38, 29.28, 29.20, 29.07, 25.81, 25.52, 14.28. HRMS (ESI-TOF) calculated for $C_{35}H_{67}NO_4S$ $[M+H]^-$ : 598.4864, found 598.4858.

Synthesis of the reactive head group **8**

Reactive head group was synthesized according to the Scheme S1.

*tert*-butyl (2-oxo-2-(((2*R*,3*R*,4*S*,5*R*,6*R*)-3,4,5-trihydroxy-6-(hydroxymethyl)tetrahydro-2*H*-pyran-2-yl)amino)ethyl)carbamate **(5)**. *ß-D-galactopyranosyl amine* (50.0 mg, 0.27 mmol, 1 equiv) was dissolved in DMF. HATU (106.32 mg, 0.27 mmol, 1 equiv), and DIPEA (122 µL, 1.39 mmol, 5.1 equiv) were added to the solution and stirred at room temperature for 10 min. Subsequently, *N*-Boc-L-Gly (49 mg, 0.405 mmol, 1.5 equiv) was added and the reaction stirred for 12 h at 37 ºC. Afterwards, the reaction was concentrated and dissolved in MeOH (500 µL), filtered using a 0.2 µm syringe-driven filter, and the product formation was confirmed by HPLC-ELSD-MS. The crude product was collected and deprotected without further purification. 32.34 mg of the crude product was collected as a white solid. HRMS (ESI-TOF) calculated for $C_{13}H_{24}N_2O_8$ $[M+Na]^+$ : 359.1425, found 359.1425.

2-amino-*N*-((2*R*,3*R*,4*S*,5*R*,6*R*)-3,4,5-trihydroxy-6-(hydroxymethyl)tetrahydro-2*H*-pyran-2-yl)acetamide **(6).** Crude product **5** was deprotected dissolving it in a 2 mL solution of TFA/$H_2O$/TES (95/2.5/2.5 % ratio) and stirring for 2 h. The formation of the deprotected product **6** was confirmed by HPLC-ELSD-MS, and the crude mixture was purified using preparative HPLC using the solvent system with *Phase A*/*Phase B* gradients [*Phase A*: $H_2O$ with 0.1% formic acid; *Phase B*: MeOH with 0.1% formic acid]. The solvent gradient ranged from 10% phase 2 to 95% phase B over 14 minutes. The product fractions were concentrated and pure product **6** was collected as a white solid and was analyzed by $H^1$ and $C^{13}$ NMR as well as HRMS (ESI-TOF) calculated for $C_8H_{16}N_2O_6$ $[M+Na]^+$ : 259.0901, found 259.0901.

*tert*-butyl ((*R*)-1-oxo-1-((2-oxo-2-(((2*R*,3*R*,4*S*,5*R*,6*R*)-3,4,5-trihydroxy-6-(hydroxymethyl)tetrahydro-2*H*-pyran-2-yl)amino)ethyl)amino)-3-(tritylthio)propan-2-yl)carbamate **(7)**. *N*-(*tert*-butoxycarbonyl)-*S*-trityl-*L*-cysteine (104 mg, 0.224 µmol, 1 equiv) was dissolved in DMF. HATU (85.5 mg, 0.224 mmol), and DIPEA (196 µL, 1.124 mmol, 5 equiv) were added to it and stirred at room temperature for 10 min. Subsequently, compound **6** (53.12 mg, 0.224 mmol, 1 equiv) was added and the reaction mixture stirred for 12 h at 37 ºC. Afterwards, DMF was removed and the crude mixture was dissolved in MeOH (500 µL), filtered using a 0.2 µm syringe-driven filter. The formation of product **7** was confirmed by HPLC-ELSD-MS, and the crude product, collected as a slightly yellow solid, was used in the next step without further purification. HRMS (ESI-TOF) calculated for $C_{35}H_{43}N_3O_9S$ $[M+Na]^+$ : 704.2612, found 704.2618.

(*R*)-2-amino-3-mercapto-*N*-(2-oxo-2-(((2*R*,3*R*,4*S*,5*R*,6*R*)-3,4,5-trihydroxy-6-(hydroxymethyl)tetrahydro-2*H*-pyran-2-yl)amino)ethyl)propenamide **(8)**. Crude product **7** was deprotected dissolving it in a 2 mL solution of TFA/$H_2O$/TES (95/2.5/2.5 % ratio) and stirring for 2 h. The reaction was then dissolved in MeOH (500 µL), filtered using a 0.2 µm syringe-driven filter, and the product formation was confirmed by HPLC-ELSD-MS. The crude product was then purified using preparative HPLC using the solvent system with *Phase A*/*Phase B* gradients [*Phase A*: $H_2O$ with 0.1% formic acid; *Phase B*: MeOH with 0.1% formic acid]. The solvent gradient ranged from 10% phase B to 95% phase B over 14 minutes. The product fractions were concentrated and analyzed by $^1H$ and $^{13}C$ NMR as well as HRMS. Pure product was collected as a white solid. $^1H$ NMR (400 MHz, $CDCl_3$): δ 3.8-4.35 (m, 6H), 3.5-3.75 (m, 6H), 3 (m, 2H), 1.4 (s, 1H).$^{13}C$ NMR (125 MHz, CDCl3): δ 170.39, 167.62, 80.27, 77.07, 74.34, 70.02, 69.10, 61.17, 42.06, 29.72, 24.96. HRMS (ESI-TOF) calculated for $C_{11}H_{21}N_3O_7S$ $[M+Na]^+$: 362.0992, found 362.0989.

Synthesis of galactolipid **9**

Galactolipid **9** was synthesized according to the Scheme S2. The pure compound was used as a reference in the HPLC-ELSD-MS analyses to confirm the formation of **9** in the chemoenzymatic synthesis.

*S*-((*R*)-3-oxo-3-((2-oxo-2-(((2*R*,3*R*,4*S*,5*R*,6*R*)-3,4,5-trihydroxy-6-(hydroxymethyl)tetrahydro-2*H*-pyran-2-yl)amino)ethyl)amino)-2-palmitamidopropyl) hexadecanethioate **(9).** A solution of compound **8**, (5.23 mg, 0.015 mmol, 1 equiv), palmitoyl-CoA (15.02 mg, 0.015 mmol, 1 equiv), and TCEP (43.19 mg, 0.075 mmol, 5 equiv) in pH 8.5 bicine buffer (3 mL) was stirred at 37 ºC for 12 h. The reaction was concentrated and then dissolved in MeOH (500 µL), filtered using a 0.2 µm syringe-driven filter, and the crude solution was purified by preparative HPLC, using the solvent system with *Phase A*/*Phase B* gradients [*Phase A*: $H_2O$ with 0.1% formic acid; *Phase B*: MeOH with 0.1% formic acid]. The solvent gradient ranged from 50% phase B to 95% phase B over 4 minutes then to 99% phase B over 13 minutes. The product fractions were concentrated and analysed by $^1H$ and $^{13}C$ NMR as well as HRMS, providing 3.21 mg (25.5%) of galactolipid **9** as a lipid film. $^1H$ NMR (400 MHz, $CDCl_3$): δ 5.65-5.75 (m, 1H), 5.4-5.6 (m, 2H), 4.8-5 (m, 1H), 3.5-4 (m, 13H), 2.7-2.8 (m, 1H), 2.5 (m, 1H), 2.2-2.35 (m, 2H), 1.26-1.33 (m, 44H), 0.88 (m, 6H). $^{13}C$ NMR (125 MHz, CDCl3): δ 190.22, 183.38, 175.50, 161.80, 104.89, 67.73, 66.81, 61.60, 54.21, 31.99, 29.37, 27.54, 25.59, 25.32, 24.85, 23.70, 23.17, 22.67, 17.77, 16.35, 13.84, 12.06. HRMS (ESI-TOF) calculated for $C_{43}H_{81}N_3O_9S$ $[M+Na]^+$: 838.5586, found 838.5583.

Phase transition temperature determination for the lipids

Phase transition temperature for both lipids **4** and **9** were determined using a Nano DSC (TA Instruments, Waters). Briefly, a 1 mL dispersion of 1 mM lipid sample was prepared in bicine buffer (pH = 8.5) and gently vortexed and degassed for 10 min before being loaded onto the DSC. The samples were screened from 25-100 ºC and a blank sample containing only buffer was used as the reference sample.

Lipid formation starting from palmitoyl-CoA

The lipid formation reaction was carried out at a 100 µL scale. In a typical reaction, 10 µL of 1 M bicine buffer, pH 8.5 (containing 100 mM TCEP), 10 µL of 20 mM reactive head group and 10 µL of 40 mM palmitoyl-CoA were added into a glass vial containing 70 µL sterile $H_2O$ and tumbled at 37 °C. In the case of lipid **4** formation, cysteine was used as the reactive head group. In the case of the galactolipid **9**, galacto-HG **8** was used as the reactive head group. The reaction was monitored over 12 h using HPLC-ELSD-MS (Figures S5, S14). For each HPLC-ELSD-MS run, a small aliquot (10 µL) of the reaction was taken and centrifuged before loading onto the column. The amount of lipid **4** generated in the chemical reaction was determined to be 1.6±0.3 mM. In the case of galactolipid **9**, 830±30 µM lipid was formed. For the quantification, an HPLC-ELSD-MS calibration curve was generated using previously synthesized lipids **4** and **9** (Figures S5c, S14c). Chromatographic separation was performed with a mobile-phase system with gradients based on *Phase A* (water + 0.1% formic acid) and *Phase B* (methanol + 0.1% formic acid). A multistep gradient at a flow rate of 1 ml/min was used with the starting condition of *Phase B* at 50%, a linear increase to 95% until 7 min, then finally to 99% until 15 min.

As a control experiment to investigate the chemoselectivity of cysteine towards palmitoyl-CoA compared to acetyl-CoA and malonyl-CoA, lipid **4** formation was set up similarly as described above, with the addition of 10 µL of 40 mM acetyl-CoA and 10 µL of 40 mM malonyl-CoA. Finally, 50 µL of sterile water was added to bring the final volume to 100 µL. The reaction mixture was tumbled at 37 °C and monitored over 12 h using HPLC-ELSD-MS (Figure S10).

Chemoenzymatic de novo membrane formation starting from acetate as carbon feedstock

Lipids were generated starting from acetate using ACS, ACC, and FAS and cysteine or compound **8** as the reactive head group. Briefly, 12.25 µL of 1 M bicine buffer, pH 8.5 (containing 100 mM TCEP), 1.75 µL of 200 mM acetate (in sterile $H_2O$), 1.75 µL of 200 mM CoASH (in sterile $H_2O$), 1.75 µL of 200 mM $HCO_3^-$ (in sterile H2O), 3.75 µL of 200 mM ATP (in sterile $H_2O$), 3.75 µL of 200 mM NADPH (in sterile $H_2O$), 5 µL of ACS (50 µM in elution buffer), 5 µL of 50 µM ACC (in sterile $H_2O$), 12.5 µL of FAS (20 µM in elution buffer), and 2.5 µL of 10 mM reactive head group were added to a glass vial and tumbled at 37 °C. The reactions were monitored over 12 h using HPLC-ELSD-MS (Figure 4A, S24C). For each HPLC-ELSD-MS run, a small aliquot (10 µL) of the reaction was taken and centrifuged to remove protein before loading onto the column. For the quantification, as described above, a calibration curve was generated using lipid standards and the same HPLC-ELSD-MS method as described above (Figures S5, S14). The retention time for the product peaks were identical to the chemically synthesized lipids **4** and **9** respectively (Figures S5, S14). The observed lipid yields were 43% for lipid **4** and 27% for lipid **9**. Small aliquots (2 µL) were taken out at various time points and placed on a glass slide for microscopy.

Chemoenzymatic de novo vesicle formation in the presence of cholesterol

A thin film of cholesterol was first prepared in a glass vial, using $N_2$ gas to dry the film. Then, the chemoenzymatic de novo lipid formation reaction was carried out as previously described in the vial containing cholesterol and tumbled at 37 °C. The final concentration of cholesterol was 10 mol% compared to the concentration of the lipid generated.

Encapsulation of HPTS in de novo vesicles

The de novo vesicle formation reactions of lipids **4** and **9** were carried out as described above. For a typical reaction, 1 µL of 1 mM HPTS solution was added to the reaction which was tumbled at 37 °C. After 12 h, product formation was confirmed by HPLC-ELSD-MS. Afterward, the reaction was transferred to a Sephadex G-50 SEC centrifuge spin column to remove the unencapsulated HPTS. The samples were centrifuged at 0.3 rcf on a benchtop centrifuge for 4 min. Finally, 1 µL of the collected fraction of vesicle solution was placed on a clean glass slide, secured by a cover slip, and imaged on a spinning disc confocal microscope (488 nm laser) to confirm the encapsulation of HPTS.

HPTS leakage assay for de novo lipid formation after melittin incubation

To validate the pore-forming capacity of melittin, HPTS leakage was monitored after its encapsulation in the de novo formed vesicles of lipids **4** and **9**. The experiments were conducted both for the vesicle formation starting from palmitoyl-CoA (chemical step) and for the chemoenzymatic de novo lipid formation reaction. Vesicle formation reactions were carried out as described above for 12 h. After confirmation of HPTS encapsulation, a microplate reader assay was used to measure the fluorescence of HPTS for each of the lipid samples. For a representative lipid, a sample was equally split in two and 10 mol% melittin (compared to **4** or **9**) was added to one of the vials. The sample was incubated for 1 h. As a negative control, the same volume of water was added to the second vial. The samples were then filtered with a Sephadex G-50 SEC column at 0.3 rcf for 4 min to remove the released HPTS. After filtration, the fluorescent output of the samples was measured again using the plate reader.

Cryo-electron microscopy protocol for vesicle formation reactions

Cryo-electron microscopy grids (Lacey Carbon Film, Electron Microscopy Sciences #LC300-Cu) were glow-discharged (Emitech K350 unit at 20 mA for 30 s), deposited with 3.5 µL of the reaction solution, blotted for 4 sec, and then plunged into liquid ethane using a Vitrobot (Mark IV, Thermo Fisher Scientific). Images were collected on a Titan Krios G3 (Thermo Fisher Scientific) operated at 300 keV equipped with a K3 detector and a 1067HD BioContinuum energy filter (Gatan) with a 15 eV slit width. Images were acquired with a total dose of 50 e/Å2 at 2.16 Å/pixel and 4 µm nominal defocus. Bilayer thickness for lipid **4** and **9** were done by radial profiling following established protocol (Figures S9, S15) (*3*).

$^{13}$C-labeled chemoenzymatic de novo lipid formation

The chemoenzymatic de novo cascade reactions were carried out as described above in the presence of HPTS. Lipid formation was monitored by HPLC-ELSD-MS for 8 h, after which, vesicle formation and HPTS encapsulation were confirmed. Subsequently, 10 mol% melittin was added to the vesicles and incubated at 37 °C for 30 min. After excess HPTS was washed off from the reaction by spin-filtration, reactive headgroup (250 µM), $^{13}$C-labeled acetate (10 mM), CoASH (10 mM), $HCO_3^-$ (7mM), ATP (15 mM), and NADPH (15 mM) were added to the respective reactions and incubated for further 8 h. For lipid **4**, cysteine was the reactive head group and for galactolipid **9** formation, galacto-HG **8** was the reactive head group. HPLC-ELSD-MS was used to quantify the amounts of $^{13}$C-labeled lipids formed.

Proton gradient assay

Lipid formation reactions starting from palmitoyl-CoA were set up as described above in the presence of HPTS in the reaction buffer for both lipid **4** and for galactolipid **9**. After 8 h, lipid formation was confirmed by HPLC-ELSD-MS and HPTS encapsulation was confirmed by confocal microscopy. Subsequently, the reaction buffer (bicine pH = 8.5) was exchanged with pH = 4.6 citrate buffer using spin-filtration and HPTS fluorescence was monitored over time using a microplate reader.

### 3. Supplementary Schemes

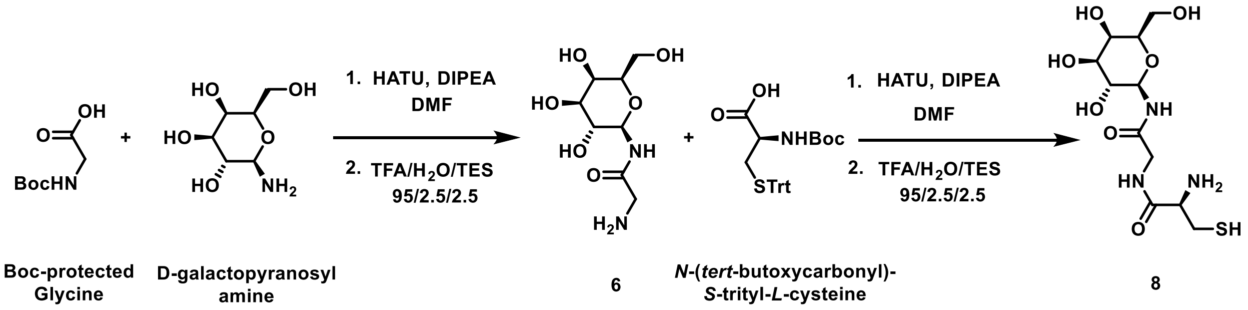


**Scheme S1.** Reaction scheme for the synthesis of head group **8**.

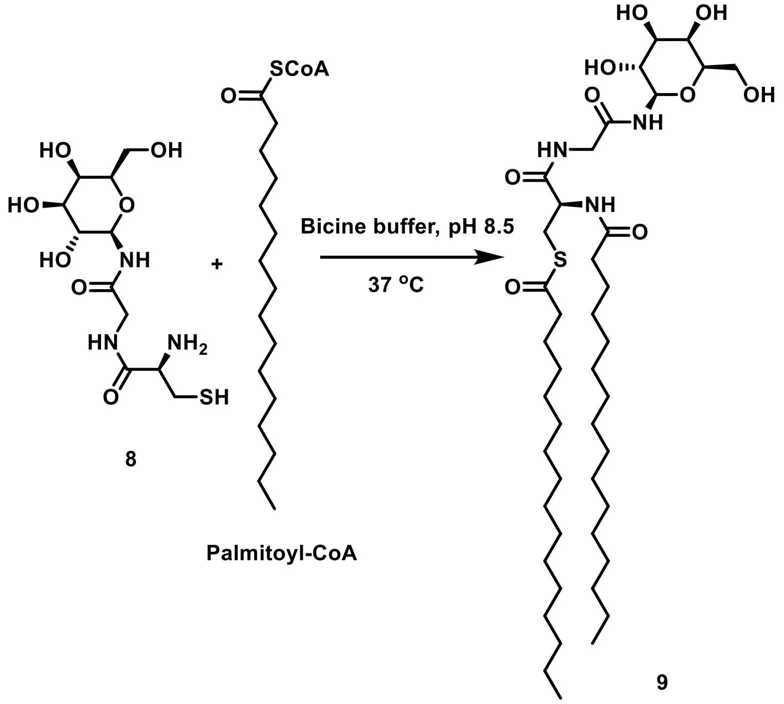


**Scheme S2.** Reaction scheme for lipid **9** synthesis from palmitoyl-CoA and galacto-HG **8** in bicine buffer pH 8.5.

### 4. Supplementary Figures

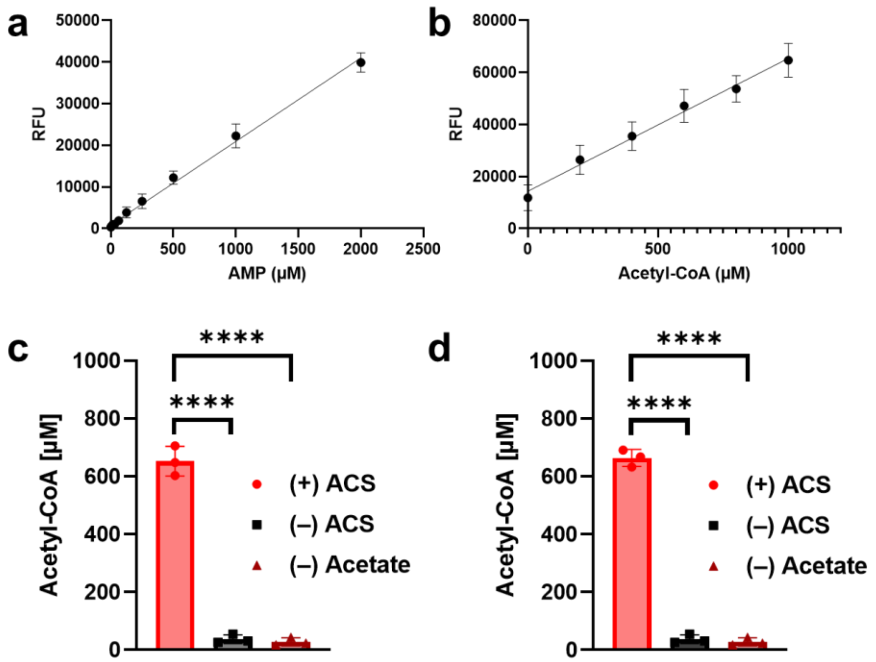


**Figure S1. (a)** Standard curve used to quantify AMP levels in experimental samples. Data points presented as means ± s.d. (*n* = 3 technically independent samples). **(b)** Standard curve used to quantify acetyl-CoA levels in experimental samples. Data points presented as means ± s.d. (*n* = 3 technically independent samples). **(c)** Activity assay of ACS determined by AMP-Glo™, indirectly showing the formation of acetyl-CoA. No acetyl-CoA is formed in the absence of acetate or ACS. Data points presented as means ± s.d. (*n* = 3 technically independent samples), and the significance determined using an unpaired t-test (two tailed). (+) ACS vs (–) ACS ****P < 0.0001; (+) ACS vs (–) acetate ****P < 0.0001. **(d)** Activity assay of ACS determined by the acetyl-CoA detection assay. No acetyl-CoA is formed in the absence of acetate or ACS. Data points presented as means ± s.d. (*n* = 3 technically independent samples), and the significance determined using an unpaired t-test (two tailed). (+) ACS vs (–) ACS ****P < 0.0001; (+) ACS vs (–) acetate ****P < 0.0001.

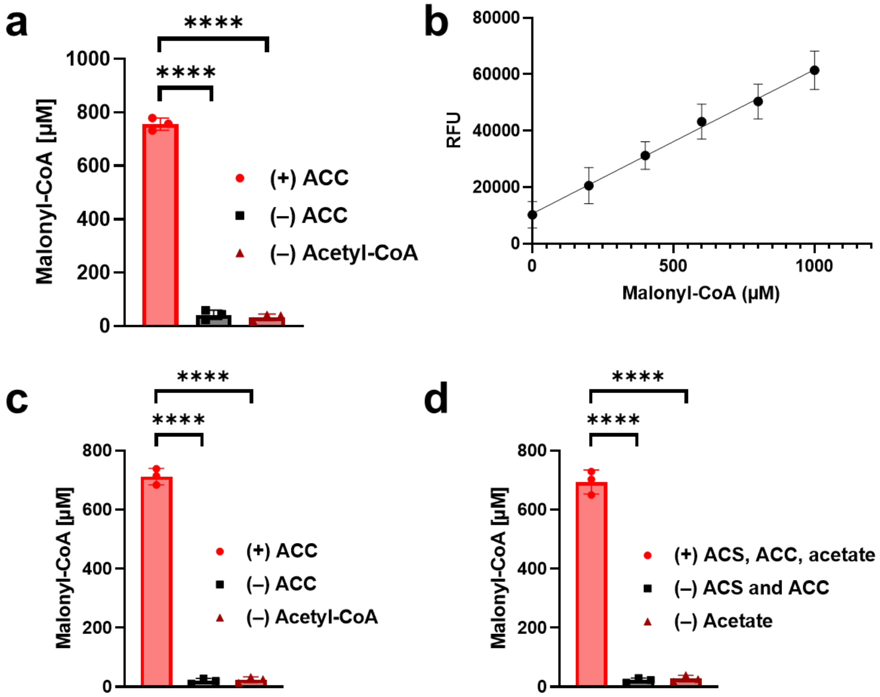


**Figure S2. (a)** Activity assay of ACC determined by AMP-Glo™, indirectly showing the formation of malonyl-CoA. No malonyl-CoA is formed in the absence of acetyl-CoA or ACC. Data points presented as means ± s.d. ($n$ = 3 technically independent samples), and the significance determined using an unpaired t-test (two tailed). (+) ACC vs (–) ACC ****P < 0.0001; (+) ACS vs (–) acetyl-CoA ****P < 0.0001. **(b)** Standard curve used to quantify malonyl-CoA levels in experimental samples. Data points presented as means ± s.d. ($n$ = 3 technically independent samples). **(c)** ACC activity assay based on malonyl-CoA quantification by ELISA. No malonyl-CoA is formed in the absence of acetyl-CoA or ACC. Data points presented as means ± s.d. ($n$ = 3 technically independent samples), and the significance determined using an unpaired t-test (two tailed). (+) ACC vs (–) ACC ****P < 0.0001; (+) ACS vs (–) acetyl-CoA ****P < 0.0001. **(d)** ACS and ACC combined activity assay, starting from acetate, based on malonyl-CoA quantification by ELISA. No malonyl-CoA is formed in the absence of acetate or enzymes. Data points presented as means ± s.d. ($n$ = 3 technically independent samples), and the significance determined using an unpaired t-test (two tailed). (+) ACS, ACC, acetate vs (–) ACS and ACC ****P < 0.0001; (+) ACS, ACC, acetate vs (–) acetate ****P < 0.0001.

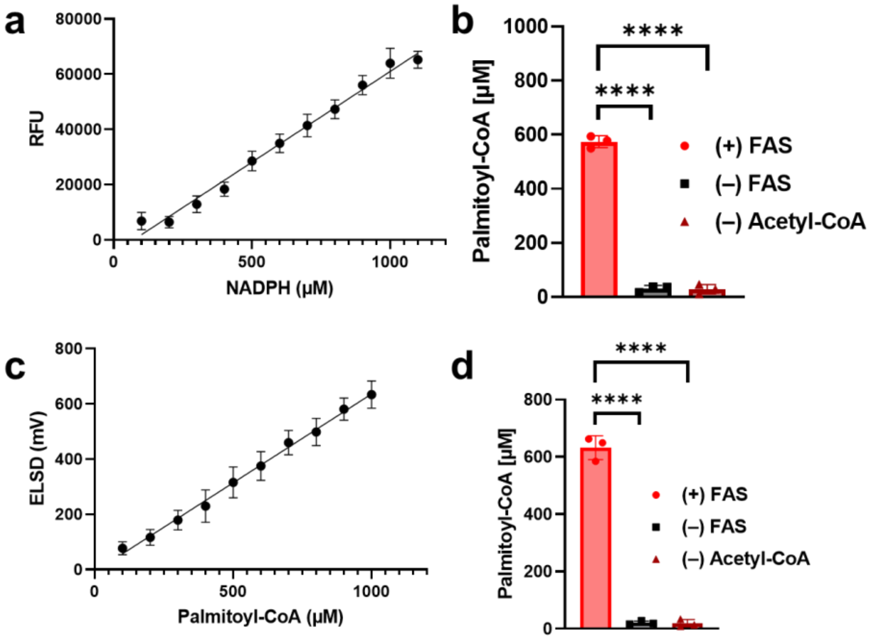


**Figure S3. (a)** Standard curve used to quantify NADPH levels in experimental samples. Data points presented as means ± s.d. ($n$ = 3 technically independent samples). **(b)** Activity assay of FAS determined by NADPH quantification, indirectly showing the formation of palmitoyl-CoA. No palmitoyl-CoA is formed in the absence of acetyl-CoA or FAS. Data points presented as means ± s.d. ($n$ = 3 technically independent samples), and the significance determined using an unpaired t-test (two tailed). (+) FAS vs (–) FAS ****P < 0.0001; (+) FAS vs (–) acetyl-CoA ****P < 0.0001. **(c)** Standard curve used to quantify palmitoyl-CoA levels in experimental samples. Data points presented as means ± s.d. ($n$ = 3 technically independent samples). **(d)** Activity assay of FAS determined by HPLC-ELSD-MS quantification of palmitoyl-CoA. No palmitoyl-CoA is formed in the absence of acetyl-CoA or FAS. Data points presented as means ± s.d. ($n$ = 3 technically independent samples), and the significance determined using an unpaired t-test (two tailed). (+) FAS vs (–) FAS ****P < 0.0001; (+) FAS vs (–) acetyl-CoA ****P < 0.0001.

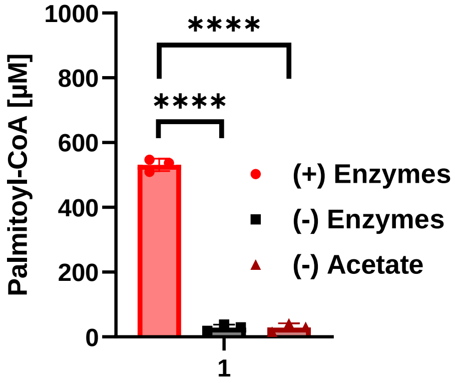


**Figure S4.** Palmitoyl-CoA synthesis from acetate mediated by the enzymatic cascade of ACS, ACC, and FAS. Data points presented as means ± s.d. ($n$ = 3 technically independent samples), and the significance determined using an unpaired t-test (two tailed). (+) Enzyme vs (–) acetate ****P < 0.0001; (+) Enzyme vs (–) Enzymes ****P < 0.0001.

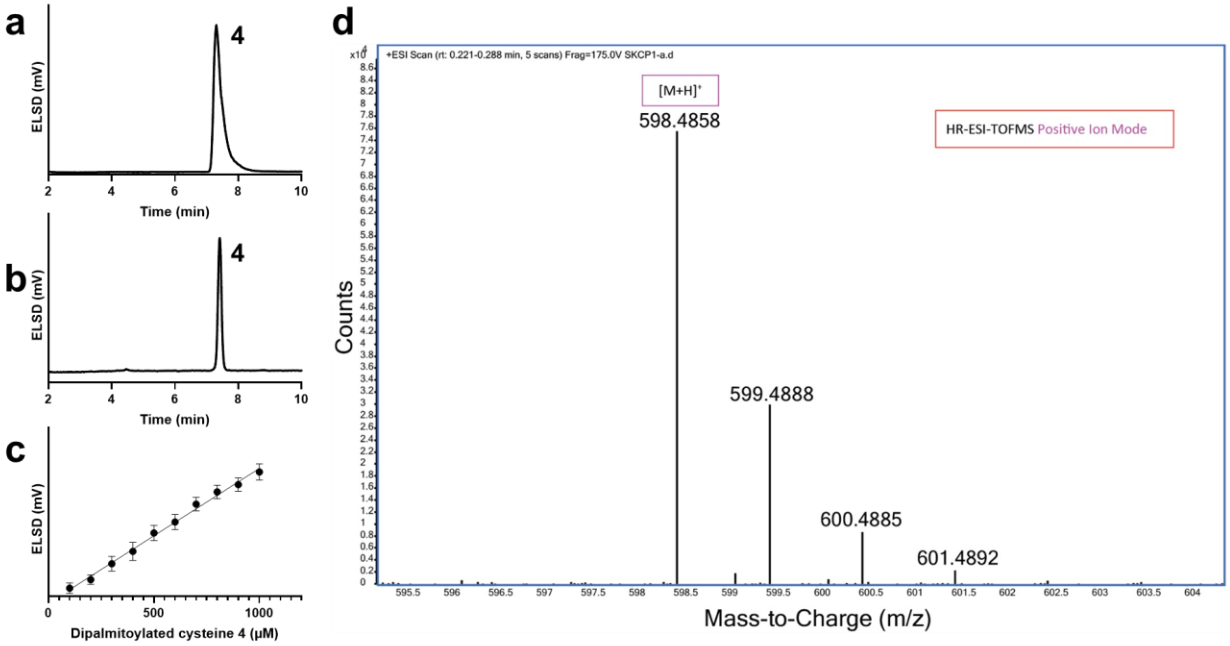


**Figure S5. (a)** HPLC-ELSD trace for the chemically synthesized dipalmitoylated cysteine **4** (standard). **(b)** HPLC-ELSD trace of dipalmitoylated cysteine **4** produced from the reaction between cysteine and palmitoyl-CoA. No monoacylated cysteine intermediate was observed during the reaction. **(c)** Standard curve used to quantify compound **4** levels in experimental samples. Data points presented as means ± s.d. ($n$ = 3 technically independent samples). **(d)** HRMS spectrum confirming the identity of compound **4**.

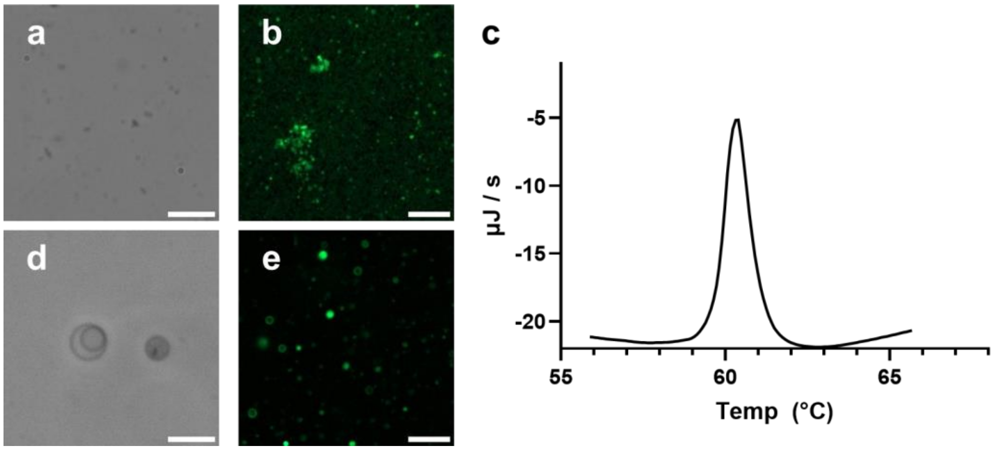


**Figure S6**. **(a)** Phase-contrast microscopy image of vesicles of lipid **4** formed by dipalmitoylation of cysteine in the absence of cholesterol, taken at 8 h reaction time. Scale bar = 10 μm. **(b)** Fluorescence microscopy image of vesicles of lipid **4** formed by dipalmitoylation of cysteine in the absence of cholesterol, taken at 8 h reaction time. Membranes were stained with 0.1 mol% BODIPY-FL. Scale bar = 10 μm. **(c)** Representative DSC trace of purified lipid **4**, indicating a phase transition of 60.4 °C. **(d)** Phase-contrast microscopy image of vesicles of lipid **4** formed by dipalmitoylation of cysteine in the presence of 10 mol% cholesterol, taken at 8 h reaction time. Scale bar = 10 μm. **(e)** Fluorescence microscopy image of vesicles of lipid **4** formed by dipalmitoylation of cysteine in the presence of 10 mol% cholesterol, taken at 8 h reaction time. Membranes were stained with 0.1 mol% BODIPY-FL. Scale bar = 10 μm.

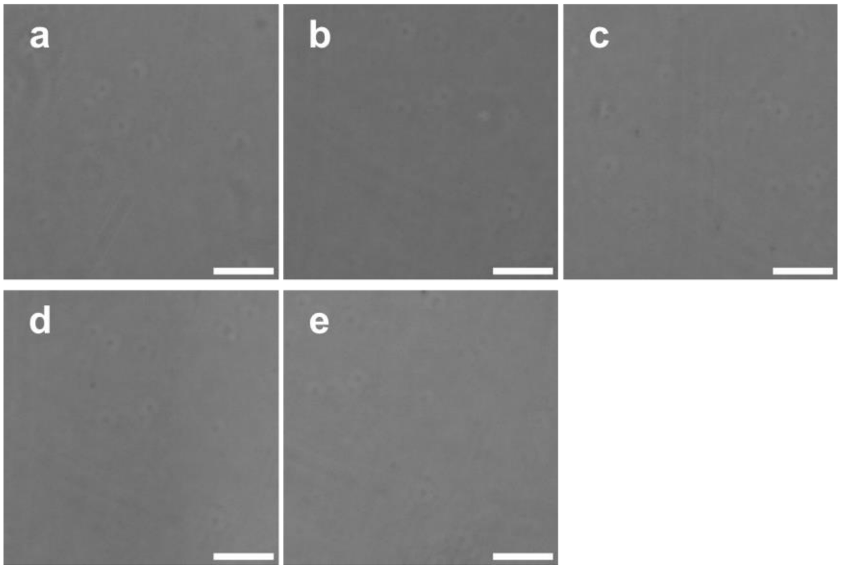


**Figure S7.** Negative controls for cholesterol addition in bicine buffer pH = 8.5. Phase-contrast microscopy image of **(a)** 50 µM cholesterol in buffer, **(b)** lipid formation reaction precursors NADPH, ATP, $HCO_3^-$ and CoASH in the presence of 50 µM cholesterol in buffer, **(c)** ACS, ACC, and FAS in the presence of 50 µM cholesterol in buffer, and **(d)** 0.5 mM cysteine in the presence of 50 µM of cholesterol and **(e)** 0.5 mM palmitoyl-CoA in the presence of 50 µM cholesterol in buffer. Scale bars = 10 µm.

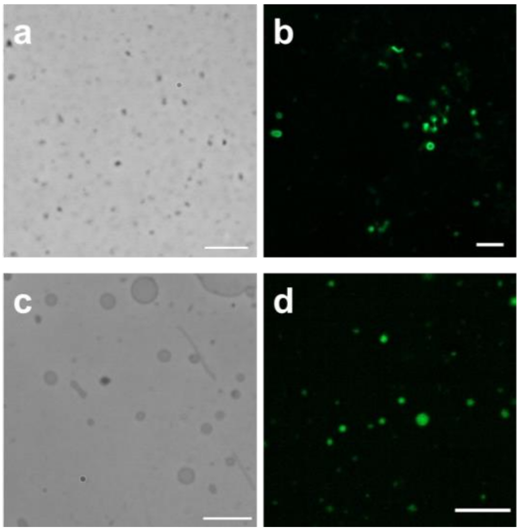


**Figure S8. (a)** Phase-contrast microscopy image of vesicles of lipid **4** generated by chemoenzymatic synthesis from acetate and cysteine as starting precursors, taken at 8 h reaction time. Scale bar = 10 µm. **(b)** Fluorescence microscopy image of vesicles of lipid **4** generated by chemoenzymatic synthesis from acetate and cysteine as starting precursors, taken at 8 h reaction time. Membranes were stained with 0.1 mol% BODIPY-FL. Scale bar = 10 µm. **(c)** Phase-contrast microscopy image of vesicles of lipid **4** generated by chemoenzymatic synthesis from acetate and cysteine as starting precursors in the presence of 10 mol% cholesterol, taken at 8 h reaction time. Scale bar = 10 µm. **(d)** Fluorescence microscopy image of vesicles of lipid **4** generated by chemoenzymatic synthesis from acetate and cysteine as starting precursors in the presence of 10 mol% cholesterol, taken at 8 h reaction time. Membranes were stained with 0.1 mol% BODIPY-FL. Scale bar = 10 µm.

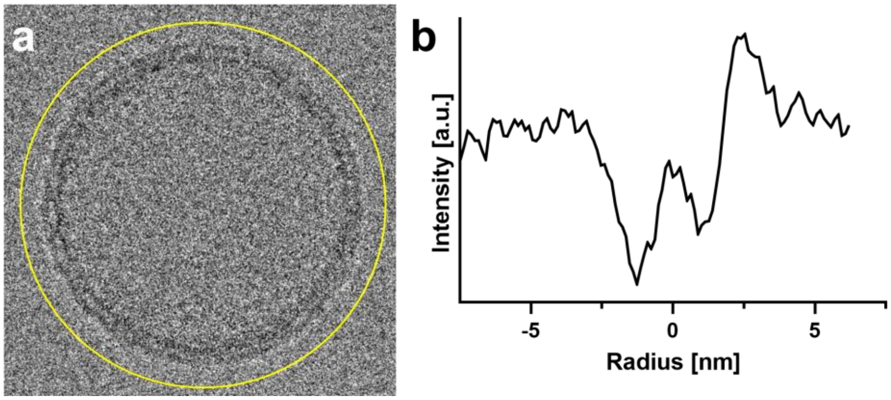


**Figure S9. (a)** Cryo-EM image of a lipid **4** vesicle generated chemoenzymatically in the presence of 10 mol% cholesterol in bicine buffer pH 8.5, and **(b)** radial intensity profile of the membrane used to estimate the bilayer thickness.

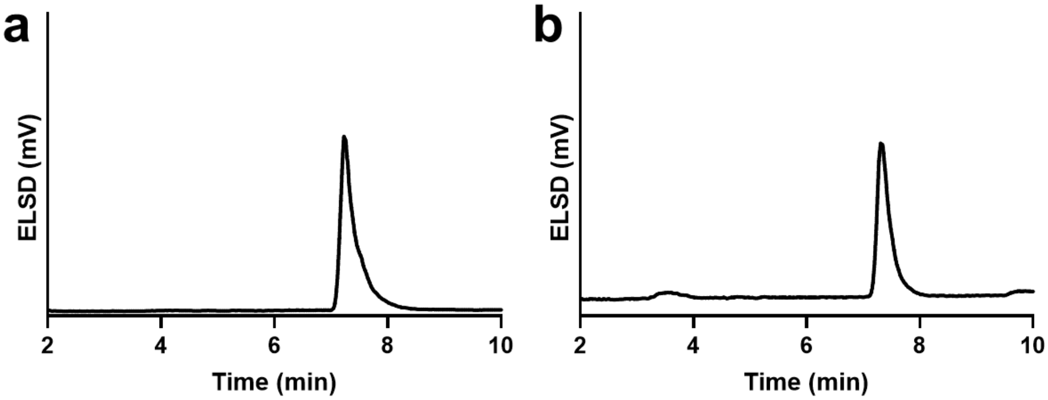


**Figure S10. (a)** HPLC-ELSD trace showing the formation of dipalmitoylated cysteine **4** by chemoenzymatic synthesis using acetate and cysteine as the reactive precursors. Data corresponds to a 12 h reaction time. **(b)** HPLC-ELSD trace showing the formation of dipalmitoylated cysteine **4** obtained from the reaction between cysteine (2 mM) and palmitoyl-CoA (4 mM) in the presence of acetyl-CoA (4 mM), malonyl-CoA (4 mM). Lipid **4** was the majority product seen after 12 h of reaction.

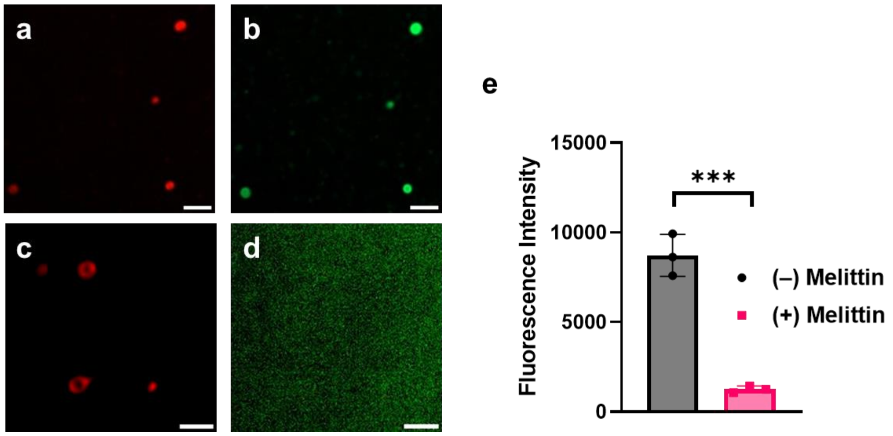


**Figure S11.** (**a-d)** Fluorescence microscopy image of lipid **4** vesicles generated by chemoenzymatic reaction, shown before **(a-b)** and after **(c-d)** incubation with 10 mol% melittin for 1 h. Membranes were stained with 0.1 mol% of Texas Red DHPE (as shown in **a** and **c**). Vesicles initially encapsulated HPTS **(b)**; following melittin treatment, HPTS leakage into the external medium was observed **(d)**. Scale bars = 10 µm. **(e)** Fluorescence intensity of vesicle samples of **4** with and without melittin showing significant leakage of HPTS upon melittin treatment. Data points presented as means ± s.d. (*n* = 3 technically independent samples), and the significance determined using an unpaired t-test (two tailed). (–) Melittin vs (+) Melittin ***P = 0.0004.

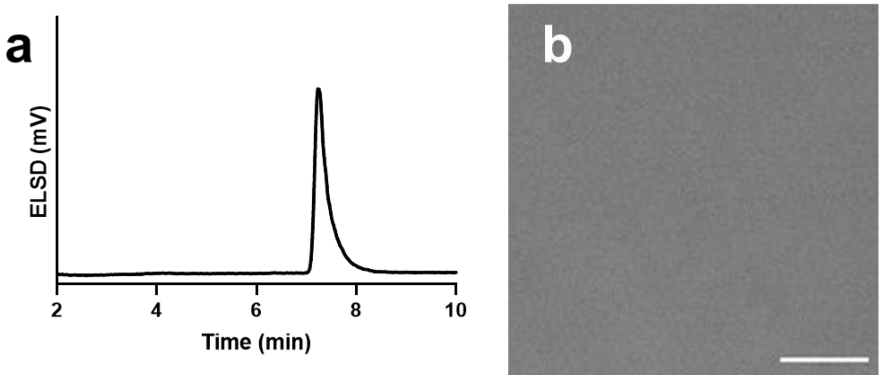


**Figure S12. (a)** HPLC-ELSD trace acquired after a 1 h incubation with 25 mol% melittin, confirming the presence of lipid **4** generated via chemoenzymatic synthesis using acetate and cysteine as precursors. (b) Phase-contrast microscopy of lipid **4** generated via chemoenzymatic synthesis acquired after a 1 h incubation with 25 mol% melittin, showing that vesicles have disassembled in the presence of excess melittin. Scale bar = 5 µm.

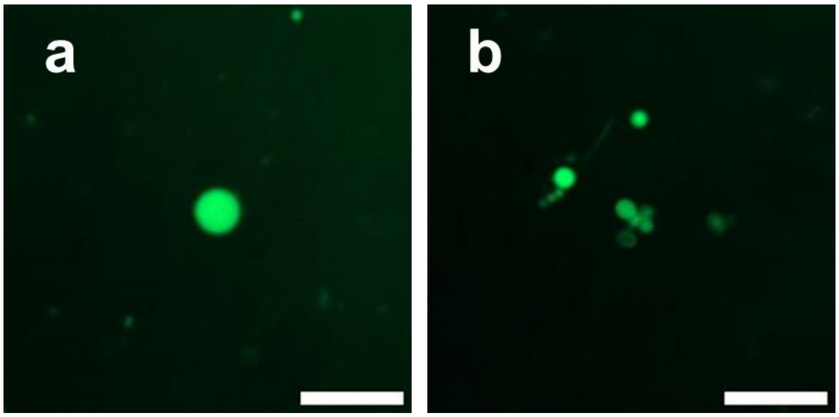


**Figure S13.** Fluorescence microscopy image of lipid **4** vesicles formed via chemoenzymatic synthesis encapsulating **(a)** HPTS (image taken after 72 h) showing that the dye is retained over extended periods when melittin is not present, and **(b)** GFP, in presence of 10 mol% melittin, showing that the protein is retained within the vesicles despite the presence of melittin. Scale bars = 10 µm.

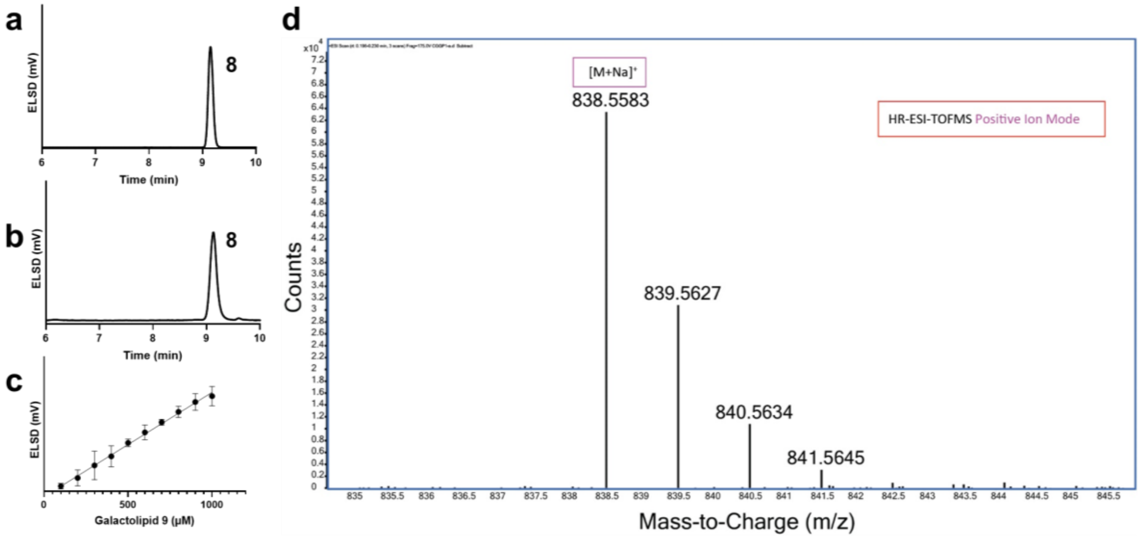


**Figure S14. (a)** HPLC-ELSD trace for the chemically synthesized compound **9** (standard). **(b)** HPLC-ELSD trace of compound **9** produced from the reaction between head group **8** and palmitoyl-CoA. **(c)** Standard curve used to quantify compound **9** levels in experimental samples. Data points presented as means ± s.d. ($n$ = 3 technically independent samples). **(d)** HRMS spectrum confirming the identity of compound **9**.

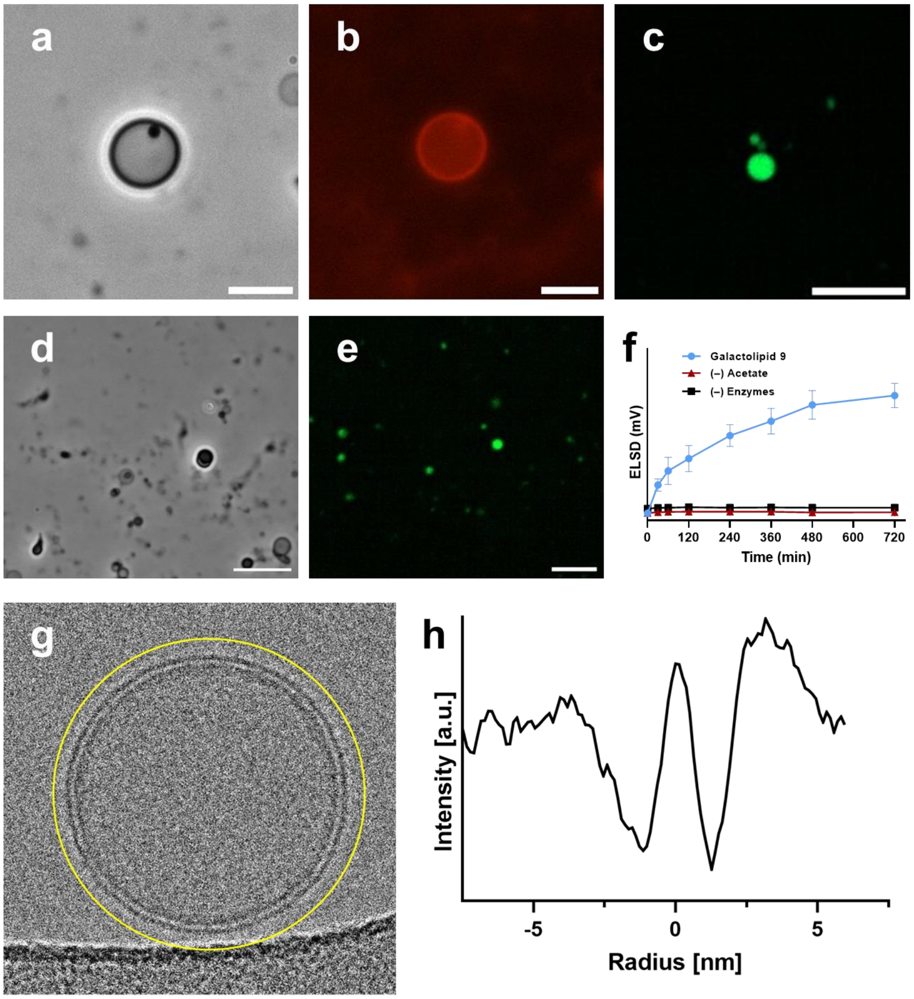


**Figure S15**. **(a)** Phase-contrast microscopy image of vesicles of galactolipid **9** formed by dipalmitoylation of galacto-HG **8** using palmitoyl-CoA in the presence of 10 mol% cholesterol, acquired at 8 h reaction time. Scale bar = 10 µm. **(b)** Fluorescence microscopy image of vesicle of galactolipid **9** formed by dipalmitoylation of galacto-HG **8** using palmitoyl-CoA in the presence of 10 mol% cholesterol, acquired at 8 h reaction time. Membranes were stained with 0.1 mol% Texas Red DHPE. Scale bar = 10 µm. **(c)** Fluorescence microscopy image showing HPTS encapsulation within the de novo formed galactolipid **9** vesicles. Scale bar = 10 µm. **(d)** Phase-contrast microscopy image of vesicles of galactolipid **9** formed via chemoenzymatic synthesis using acetate and cysteine as starting precursors, in the presence of 10 mol% cholesterol. Image acquired at 8 h reaction time. **(e)** Fluorescence microscopy image showing HPTS encapsulation within vesicles of galactolipid **9** formed via chemoenzymatic synthesis using acetate and cysteine as starting precursors, in the presence of 10 mol% cholesterol. Image acquired at 8 h reaction time. Scale bars = 10 µm. **(f)** Kinetic curve corresponding to the chemoenzymatic generation of galactolipid **9** over 12 h. Negative controls show that no product was formed in the absence of acetate or ACS, ACC, and FAS enzymes. Data points presented as means ± s.d. (*n* = 3 technically independent samples). **(g)** Cryo-EM image of a lipid **9** vesicle generated via chemoenzymatic synthesis in the presence of 10 mol% cholesterol in bicine buffer pH 8.5, and **(h)** radial intensity profile of the membrane used to estimate the bilayer thickness.

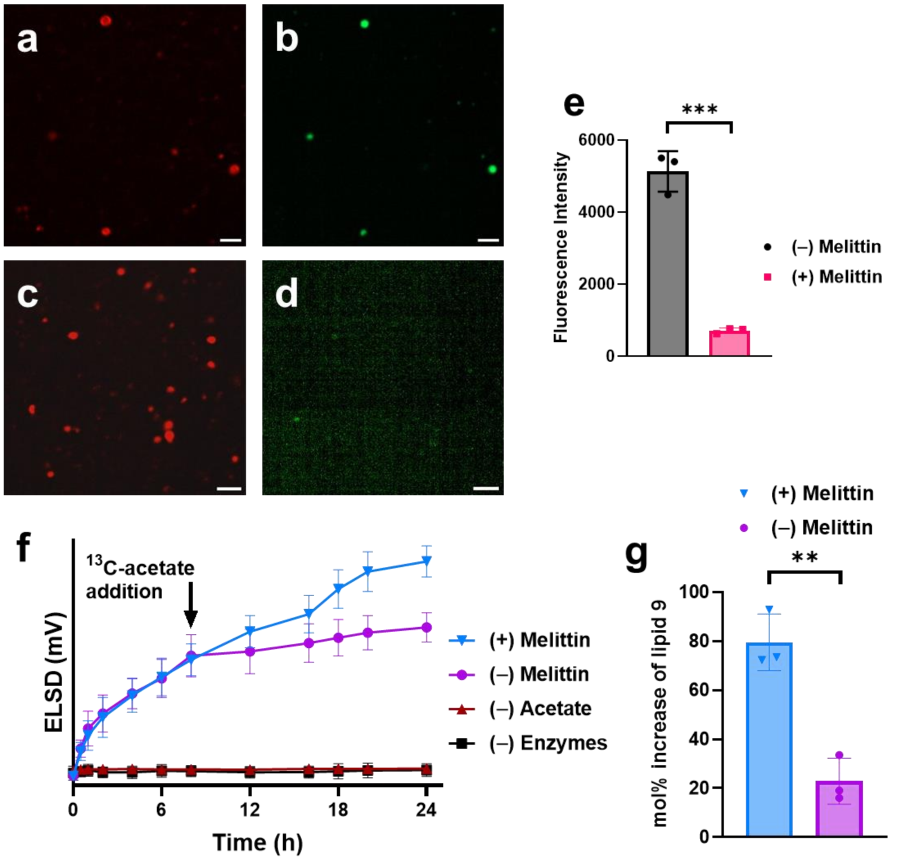


**Figure S16.** **(a-d)** Fluorescence microscopy image of lipid **9** vesicles generated by chemoenzymatic synthesis, shown before **(a-b)** and after **(c-d)** incubation with 10 mol% melittin for 1 h. Membranes were stained with 0.1 mol% of Texas Red DHPE (as shown in **a** and **c**). Vesicles initially encapsulated HPTS **(b)**; following melittin treatment, HPTS leakage into the external medium was observed **(d)**. Scale bars = 10 µm. **(e)** Fluorescence intensity of vesicle samples of **9** with and without melittin showing significant leakage of HPTS upon melittin treatment. Data points presented as means ± s.d. (*n* = 3 technically independent samples), and the significance determined using an unpaired t-test (two tailed). (–) Melittin vs (+) Melittin ***P = 0.0002. **(f)** Chemoenzymatic de novo formation of lipid **9** monitored over time. 10 mol% melittin was added after 8 h of diacylation and incubated for 1 h. Fresh reaction precursors including $^{13}$C-acetate and compound **8** were added and the reaction followed for a further 8 h. As shown in the graph, significant additional $^{13}$C-labeled lipid **9** is synthesized when melittin is present (79% increase). Negative control without melittin addition showed reduced formation of lipid **9** (23% increase). No lipid formation was observed in controls lacking acetate or enzymes. Data points presented as means ± s.d. (*n* = 3 technically independent samples). **(g)** Increase in the amount of lipid **9** after the addition of new reaction precursors in the presence (79% increase) and absence of melittin (23% increase). Data points presented as means ± s.d. (*n* = 3 technically independent samples), and the significance determined using an unpaired t-test (two tailed). (–) Melittin vs (+) Melittin **P = 0.0027.

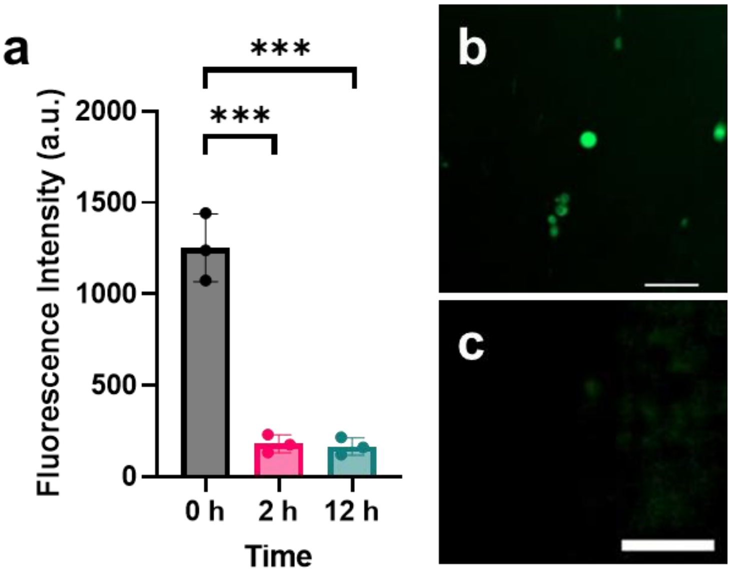


**Figure S17.** Proton gradient assay for oleic acid vesicles. A thin film of oleic acid was prepared and then rehydrated in bicine buffer (pH = 8.5) containing 0.1 mol% HPTS. The vesicles were tumbled for 1 h. **(a)** Fluorescence intensity was measured at 0 h, 2 h, and 12 h after buffer exchange from bicine to citrate (pH = 4.6). Data points presented as means ± s.d. (*n* = 3 technically independent samples), and the significance determined using an unpaired t-test (two tailed). 0 h vs 2 h ***P = 0.0006; 0 h vs 12 h ***P = 0.0006. **(b-c)** Fluorescence microscopy images before **(b)** and after **(c)** buffer exchange.

### 5. Characterization Spectra

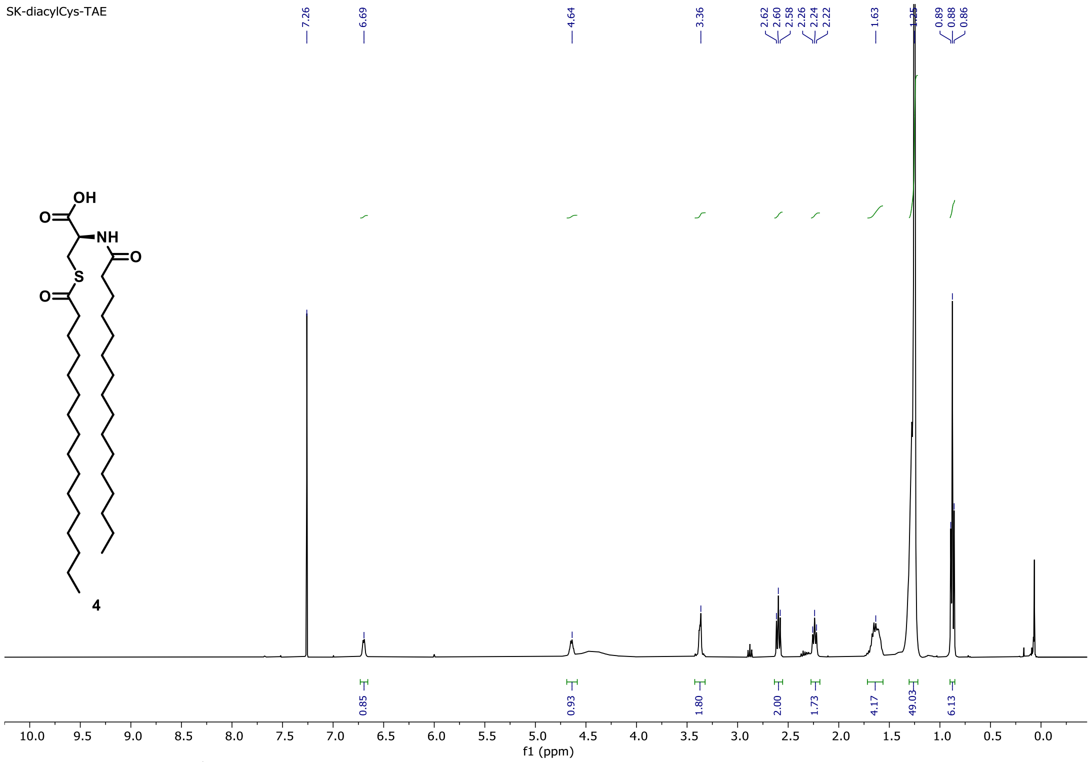


$^{1}H$ NMR spectrum of Dipalmitoylated Cysteine **4** in $CDCl_3$

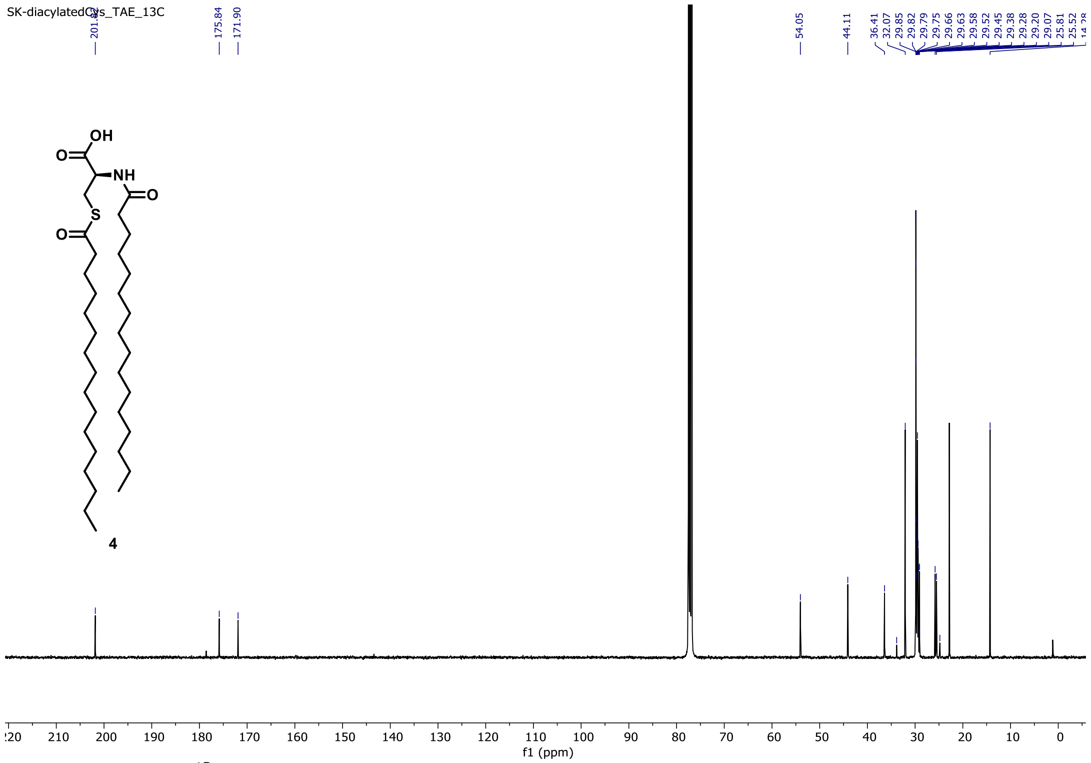


$^{13}C$ NMR spectrum of Dipalmitoylated Cysteine **4** in $CDCl_3$

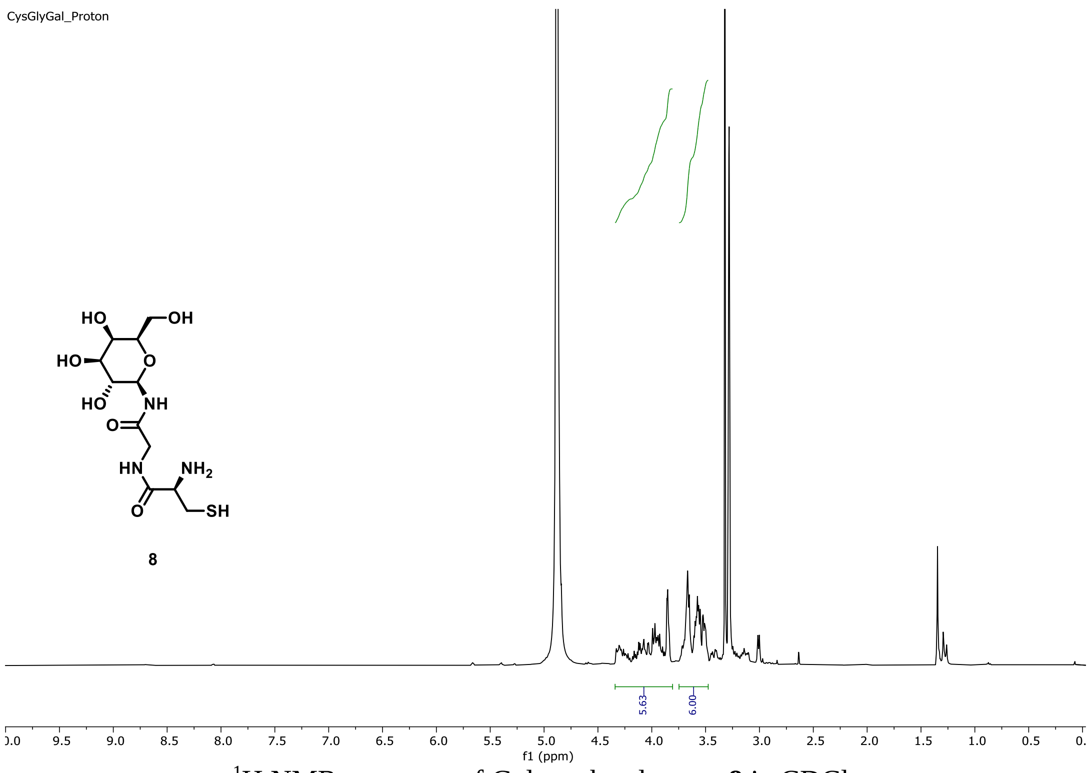


$^{1}$H NMR spectrum of Galacto head group **8** in $CDCl_3$

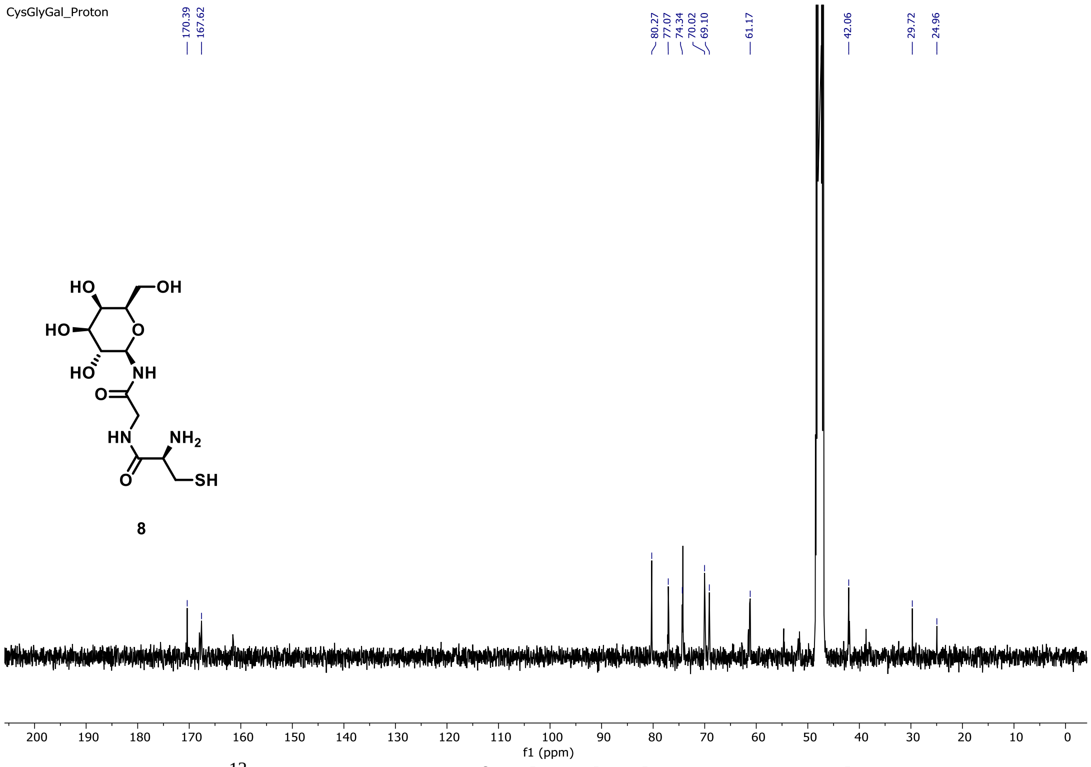


$^{13}$C NMR spectrum of Galacto head group **8** in $CDCl_3$

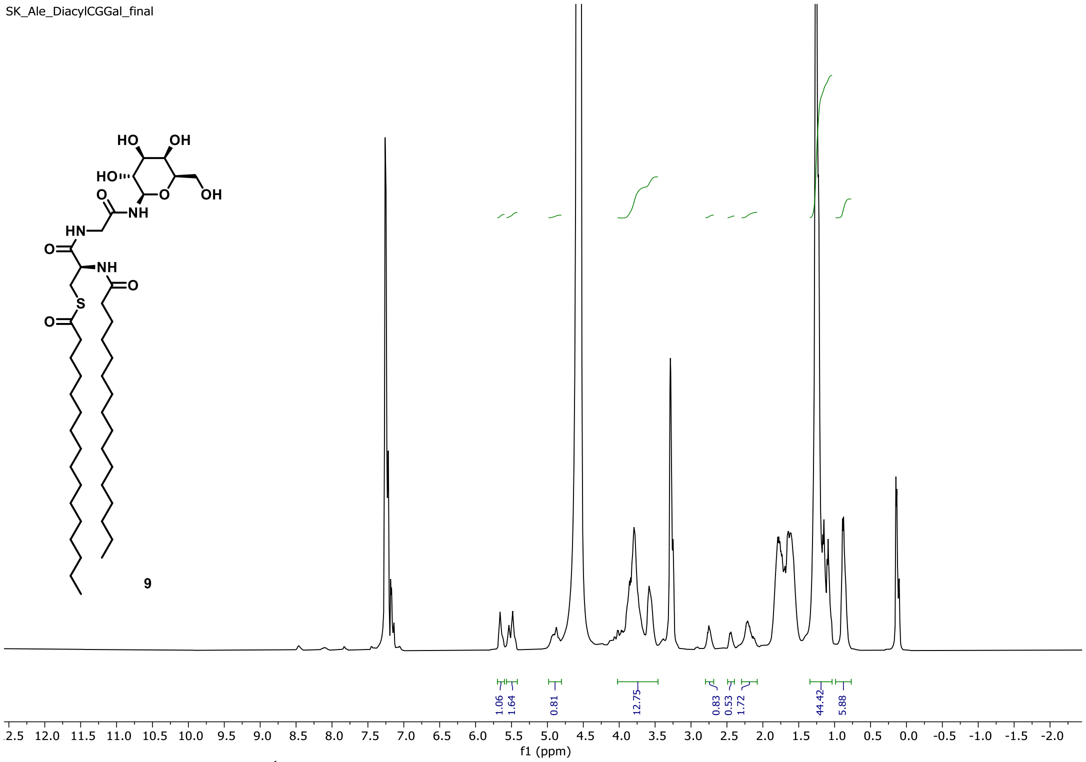


$^1$H NMR spectrum of Galactolipid **9** in $CDCl_3$.

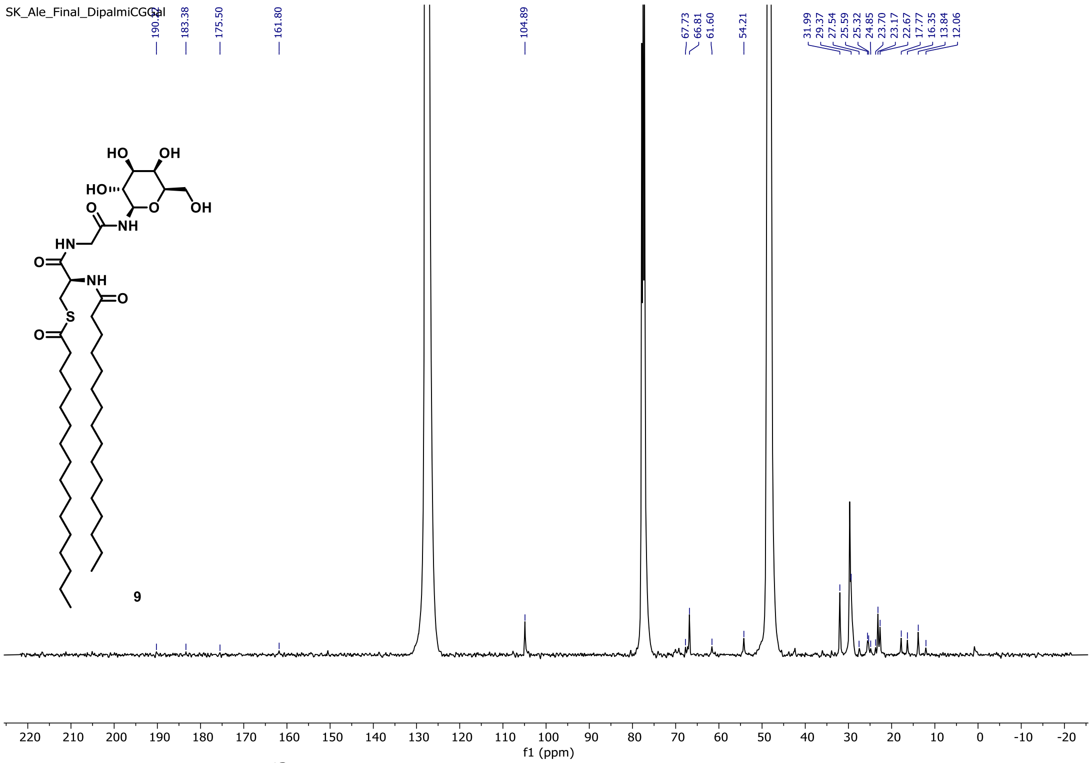


$^{13}$C NMR spectrum of Galactolipid **9** in $CDCl_3$.
